# An Inducible *Cre* Mouse with Preferential Activity in Vascular Smooth Muscle Evades a Previously Lethal Intestinal Phenotype

**DOI:** 10.1101/2022.02.03.479061

**Authors:** Ganesh D. Warthi, Jessica L. Faulkner, Jaser Doja, Amr R. Ghanam, Pan Gao, Allison C. Yang, Orazio J. Slivano, Candee T. Barris, Taylor C. Kress, Scott D. Zawieja, Susan H. Griffin, Xiaoling Xie, Alan Ashworth, Christine K. Christie, William B. Bryant, Ajay Kumar, Michael J. Davis, Xiaochun Long, Lin Gan, Eric J. Belin de Chantemèle, Qing Lyu, Joseph M. Miano

## Abstract

All smooth muscle cell (SMC) restricted *Cre* mice recombine floxed alleles in vascular and visceral SMCs. We generated a new tamoxifen-inducible *CreER^T2^* mouse, *Itga8-CreER^T2^*, and compared its activity to the widely used *Myh11-CreER^T2^* mouse. Both *CreER^T2^* mice showed similar activity in vascular SMCs; however, *Itga8-CreER^T2^* displayed limited activity in visceral SMC-containing tissues (*e.g.*, intestine). *Myh11-CreER^T2^* (but not *Itga8-CreER^T2^*) mice displayed high levels of CreER^T2^ protein, tamoxifen-independent activity, and an altered transcriptome. Whereas *Myh11-CreER^T2^*-mediated knockout of *Srf* resulted in a lethal intestinal phenotype, loss of *Srf* with *Itga8-CreER^T2^* (*Srf^Itga8^*) revealed viable mice with attenuated vascular SMC contractile gene expression, but no evidence of intestinal pathology. Male and female *Srf^Itga8^* mice presented with vascular contractile incompetence; however, only male *Srf^Itga8^* mice showed systemic changes in blood pressure. These results establish the *Itga8-CreER^T2^* mouse as an alternative to existing SMC *Cre* strains, including *Myh11-CreER^T2^*, where SMC gene loss results in visceral myopathies that obfuscate accurate phenotyping in vascular SMCs.

## Introduction

Conditional inactivation of a gene requires two components derived from bacteriophage P1: a *<u>C</u>*yclization *<u>re</u>*combination (*Cre*) gene encoding a 343 amino acid (∼38 kDa) tyrosine site-specific recombinase and 34-base pair *<u>lo</u>*cus of crossing (*<u>x</u>*) over of <u>P</u>1 (*lox*P) sequences having dyad symmetry, flanking (or floxing) the region of DNA to be excised.^1, 2^ The first application of the *Cre*/*lox*P system for conditional knockout of a gene in mice utilized an *Lck* promoter-driven *Cre* transgene to delete the 5’ promoter and first exon of a floxed *Polb* gene in T cells.^3^ This pioneering study underscores two critical issues in the design of conditional knockout mouse experiments, namely judicious placement of *lox*P sites around a sequence to be excised and restricted spatio-temporal expression of *Cre*.^4^ The spatial control of *Cre* expression is of utmost significance for the inactivation of genes in smooth muscle cells (SMCs) which are pervasive across the body plan, investing blood vessels and all hollow organs of the abdominal cavity.

SMCs have distinct embryological origins^5, 6^ and contractile properties^7–9^, but they express a common set of cell-restricted genes that maintain normal SMC homeostasis.^10^ Several of these genes have been harnessed for the development of SMC *Cre* driver mice, each of which has advantages and disadvantages.^11^ A shared characteristic of all current SMC *Cre* driver mice is the recombination activity in both vascular and visceral SMC lineages.^11^ SMC-wide Cre activity can lead to severe, often lethal, visceral myopathies^12–17^ that may impede efforts to understand the biology of key genes within vascular SMCs (VSMCs). Moreover, whole body knockout of SMC-restricted genes each resulted in embryonic or postnatal death.^18–21^ Various gene editing strategies will accelerate new floxed genes for conditional knockout studies in SMCs^22–25^, and it is likely many gene products will be of vital importance for normal visceral SMC homeostasis. Clearly, there will be instances where a more restrictive VSMC *Cre* driver mouse will be necessary to circumvent the confounding effects of visceral myopathies.

Alpha 8 integrin (*Itga8*) was first cloned from an embryonic chicken cDNA library and the ∼160 kDa protein product was shown to interact with the beta 1 integrin subunit.^26^ Independent studies reported ITGA8 protein in VSMCs, intestinal SMCs, and mesangial cells (a SMC-like cell) of the adult rat kidney.^27, 28^ A screen for retinoid-inducible genes in VSMCs reported a partial cDNA to rat *Itga8* that was used to show prominent *Itga8* mRNA expression in the adult rat aorta, with no detectable signal in a panel of other adult tissues.^29^ Subsequent studies demonstrated the down-regulation of ITGA8 with vascular injury^30^ and its role in the maintenance of VSMC differentiation.^31^ Interestingly, the Myocardin (MYOCD) coactivator^32^ stimulated *Itga8* expression in a serum response factor (SRF)-independent manner.^33^ The latter finding distinguishes *Itga8* from most other SMC markers that have functional SRF-binding sites in the promoter or first intron.^34^

A previous report found *Itga8* mRNA to be expressed abundantly in VSMCs with little to no detectable expression in most visceral SMC-containing tissues of mouse, rat, and human origin. This restricted pattern of expression in VSMCs across multiple species prompted the suggestion that the *Itga8* gene might represent an ideal locus for targeting with an inducible *Cre* in order to establish a new mouse model for more restrictive conditional knockout studies in VSMCs.^33^ Here, the generation and characterization of an *Itga8-CreER^T2^* mouse is reported. Comparative studies indicate important advantages of the *Itga8-CreER^T2^* mouse over all other SMC *Cre* drivers for the unequivocal determination of gene function in VSMCs.

## Results

### *Itga8-CreER^T2^* mice express normal levels of ITGA8 protein and a nuclear LncRNA

Among known SMC markers, the *Itga8* gene has the distinction of being expressed preferentially in VSMCs (Supplementary Fig. 1). We therefore targeted the first exon of *Itga8* with a *CreER^T2^* cassette^35^ in embryonic stem cells to establish a new inducible *Cre* driver mouse (Supplementary Fig. 2). Mice homozygous null for *Itga8* display renal agenesis and postnatal death (Supplementary Fig. 3).^36^ Because ITGA8 protein expression was not reported in heterozygous knockout mice,^36^ it was important to determine whether one functional *Itga8* allele is sufficient for normal levels of ITGA8 protein expression. Results showed no change in ITGA8 protein, but a small reduction of *Itga8* mRNA in the aorta of heterozygous *Itga8-CreER^T2^* mice (Supplementary Fig. 4). Further, there was little change in expression of a 5’ overlapping nuclear long non-coding RNA (Supplementary Fig. 4). Male and female *Itga8-CreER^T2^* mice displayed similar levels of ITGA8 and CreER^T2^ protein in aorta (Supplementary Figure 4) and they bred well with no overt signs of pathology. These results suggest a compensatory increase in ITGA8 expression from the remaining functional *Itga8* allele, supporting accurate phenotyping of conditional VSMC knockout phenotypes without the potential confounding effects of reduced ITGA8 protein.

### *Itga8-CreER^T2^* mice display preferential activity in VSMCs

More than 300 floxed genes have been targeted with various SMC *Cre* driver mice, 90% of which were generated with several versions of *Sm22-Cre* or *Myh11-Cre* (Supplementary Table 1). Previous work with *Tagln-Cre* (aka *Sm22-Cre*) reported recombination in myeloid cell lineages.^37^ To further address myeloid cell activity in SMC *Cre* mice, circulating cells from male and female *Itga8-CreER^T2^* mice carrying the *mT/mG* reporter were isolated and compared to similarly prepared cells from male *Sm22-Cre*^38^ and *Myh11-CreER^T239^* mice. As expected, fluorescence-activated cell sorting revealed ∼22% GFP positive cells in *Sm22-Cre* mice; however, no GFP fluorescence was observed in circulating cells of *Itga8-CreER^T2^* or *Myh11-CreER^T2^* mice suggesting the absence of CreER^T2^ activity in myeloid cells of these two SMC *Cre* drivers (Supplementary Fig. 5). Given the broader activity of *Sm22-Cre*,^37, 38^ the balance of experiments were conducted only in *Itga8-CreER^T2^* and *Myh11-CreER^T2^* mice.

Confocal immunofluorescence microscopy was used to compare the activity of *Itga8-CreER^T2^* with *Myh11-CreER^T2^* across multiple tissues from age-matched mice carrying the same *mT/mG* reporter used in the above flow studies. The absence of CreER^T2^ activity is indicated by red (tomato) fluorescence whereas CreER^T2^-mediated excision of a stop floxed cassette unveils green (GFP) fluorescence. Both *Itga8-CreER^T2^* and *Myh11-CreER^T2^* showed activity in SMCs of the aorta, mesentery, and microvasculature of brain and visceral organs such as intestine and bladder (Fig. 1). Higher magnification, z-stacked confocal images revealed obvious GFP fluorescence in SMCs of the aorta and vena cava with no observable GFP in endothelial cells or adventitial cells (Supplementary Fig. 6). *Myh11-CreER^T2^* showed high activity in visceral SMCs of bladder, colon, esophagus, intestine, stomach, and ureter (Fig. 1). In contrast, male and female *Itga8-CreER^T2^* mice displayed lower activity in each of these visceral SMC-enriched tissues (Fig. 1). Low *Itga8-CreER^T2^* activity was also seen in uterus (Fig. 1); no comparison could be made in uterus with *Myh11-CreER^T2^* because this transgene integrated on the Y chromosome (see below).^39^ There was little difference in activity between the two *Cre* drivers in tracheal and bronchiolar SMCs of the respiratory system (Fig. 1). Neither *Itga8-CreER^T2^* nor *Myh11-CreER^T2^* showed detectable activity (outside of blood vessels) in tissues of the eye, heart, liver, pancreas, skeletal muscle, and spleen as well as germ cells of the testis and ovary (Supplementary Fig. 6). However, *Myh11-CreER^T2^* mice exhibited pronounced activity in cells of the thymus and *Itga8-CreER^T2^* mice disclosed activity in glomerular cells of the kidney (Supplementary Fig. 6). These results demonstrate some similarities, but important differences, in the activity of *Myh11-CreER^T2^* and *Itga8-CreER^T2^*.

**Fig. 1.**
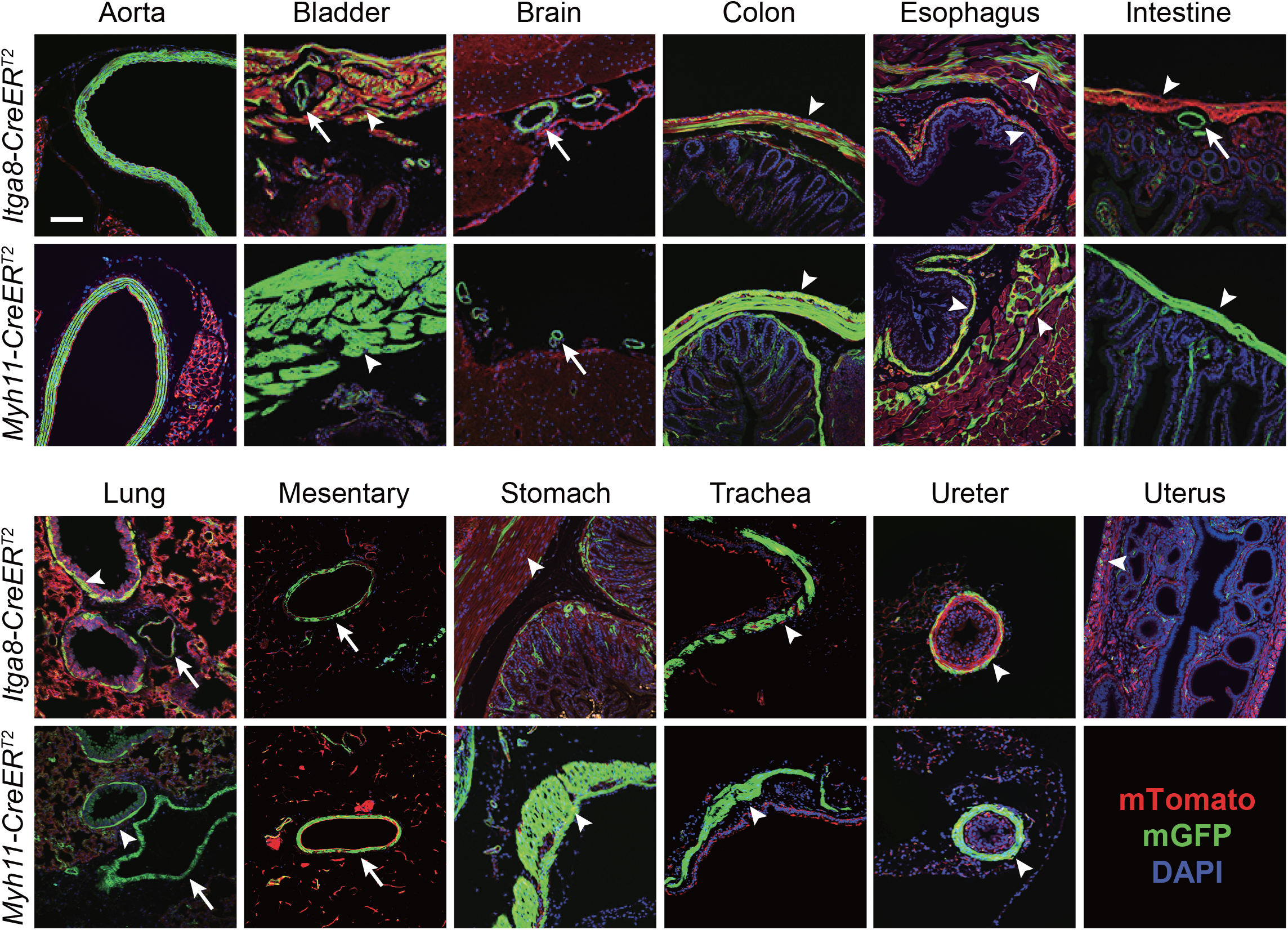
Comparative activity of *Itga8-CreER^T2^* versus *Myh11-CreER^T2^* in adult tissues. Paired tissues from 8-week old mice carrying indicated *CreER^T2^* and *mT/mG* reporter. Images shown here and below are representative of tissues from both genders (*Itga8-CreER^T2^* only) obtained over a span of three years and multiple generations. Green fluorescent protein (GFP) staining reflects Cre-mediated excision of the membrane Tomato (*mT*) reporter and is primarily confined to SMCs of each tissue. DAPI was applied where shown for nuclear staining. White arrows indicate blood vessels and white arrowheads indicate visceral smooth muscle cells. The scale bar represents 50μm for aorta and uterus and 100μm for all other panels.

### *Myh11-CreER^T2^* mice exhibit leaky activity and an altered basal transcriptome

There is evidence for tamoxifen-independent (leaky) recombination events with the *CreER^T2^* cassette.^40^ Accordingly, individually-housed male *Itga8-CreER^T2^* or *Myh11-CreER^T2^* mice carrying the *mT/mG* reporter were analyzed for unscheduled recombination. Whereas no detectable recombination of the *mT/mG* reporter was evident in *Itga8-CreER^T2^* mice, clear evidence of tamoxifen-independent recombination (ie, GFP fluorescence) was apparent in multiple tissues of *Myh11-CreER^T2^* mice (Fig. 2a). To begin to explore the basis for such leaky activity, levels of CreER^T2^ protein were analyzed in the aorta and bladder of age-matched male mice. Western blotting showed much higher CreER^T2^ protein expression in the aorta and bladder of *Myh11-CreER^T2^* mice than *Itga8-CreER^T2^* mice (Fig. 2b). Sanger sequencing disclosed the *Cre* in *Myh11-CreER^T2^* to be codon optimized for more efficient translation of the bacterial Cre protein in mammals (Supplementary Fig. 7).^41^ Further, quantitative PCR of genomic DNA revealed two copies of the *Myh11-CreER^T2^* transgene had integrated in the genome (Fig. 2c). Notably, long read sequencing demonstrated X chromosome-specific sequence near the pseudoautosomal region (PAR), flanking the integrated *Myh11-CreER^T2^* transgenes, suggesting an initial integration on the X chromosome followed by translocation to the Y chromosome (Fig. 2d). Bulk RNA-seq studies disclosed elevated expression of 101 genes in the aorta of *Myh11-CreER^T2^* mice as compared to wild type and *Itga8-CreER^T2^* mice (Fig. 2e), including two X-linked genes (*Arhgap6* and *Hccs*) found in close proximity to the *Myh11-CreER^T2^* transgenes (Fig. 2d, 2e). Collectively, these results demonstrate high level CreER^T2^ protein expression and an unusual chromosomal translocation event in *Myh11-CreER^T2^* mice, each of which appears to contribute to tamoxifen-independent activity and an altered baseline transcriptome.

**Fig. 2.**
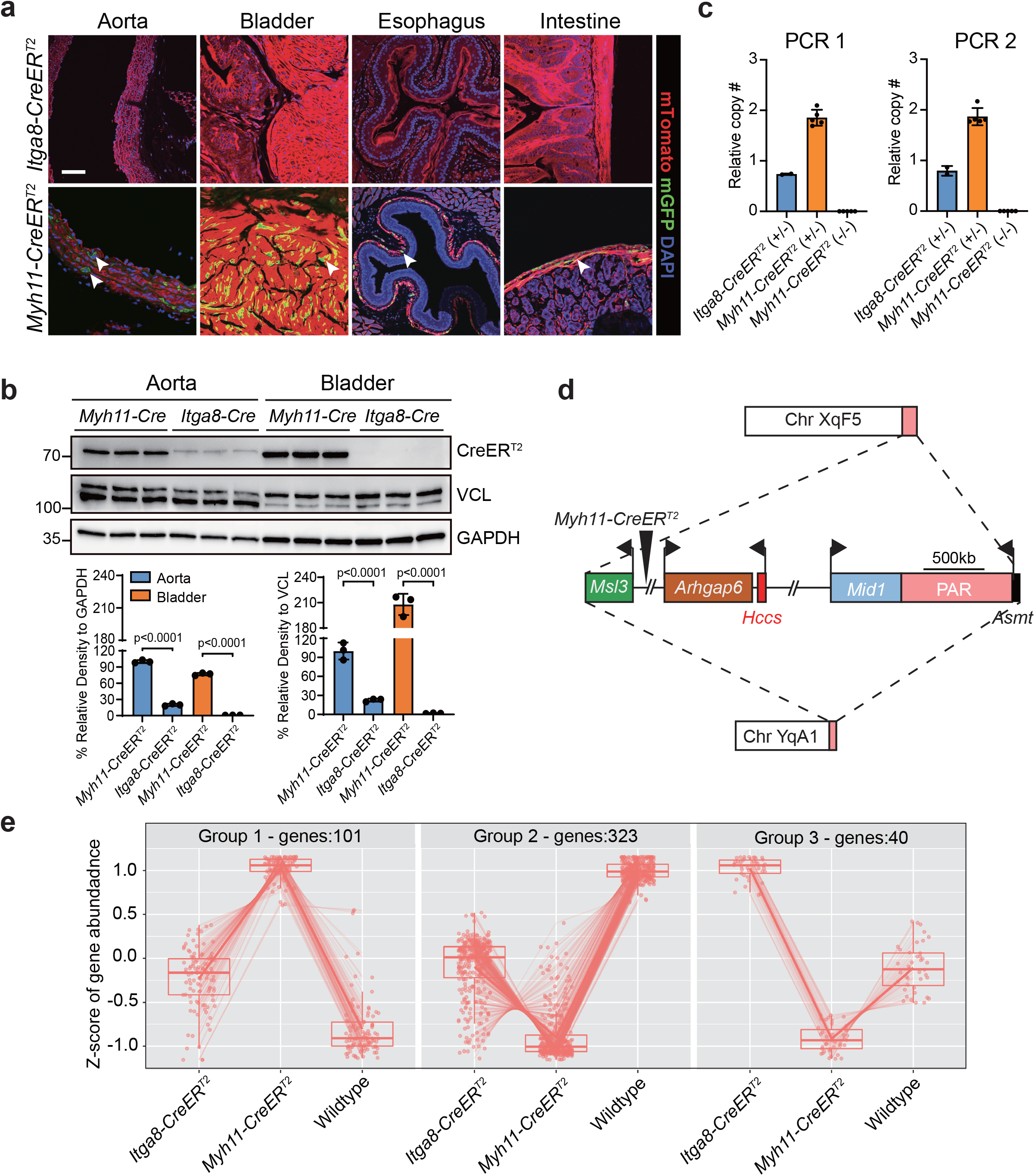
Distinguishing features of *Myh11-CreER^T2^*. **a,** Tamoxifen-independent recombination activity in *Myh11-CreER^T2^* adult tissues. Arrowheads point to SMCs with leaky activity. **b,** Western blot of *CreER^T2^* protein with quantitation below. **c,** qPCR of copy number of *Myh11-CreER^T2^* transgene from adult spleen. Two primer pairs (PCR 1 and PCR 2) were used that amplify non-overlapping sequences in *ER^T2^*. **d,** Schematic based on long read sequence mapping of *Myh11-CreER^T2^* transgene. The dashed lines indicate translocation from the X chromosome to Y chromosome carrying X-linked genes surrounding the approximate insertion site of the transgene. Bent arrows indicate the transcription start site of each gene. Scale bar is an approximation. **e,** Bulk RNA-seq summary of differentially expressed genes in the absence of tamoxifen between wild type, *Myh11-CreER^T2^* and *Itga8-CreER^T2^* mice (n = 3 mice per genotype). Group 1 represents significantly elevated genes in *Myh11-CreER^T2^* aorta as compared to *Itga8-CreER^T2^* and wild type; Group 2 represents significantly elevated genes in wild type aorta; Group 3 represents significantly elevated genes in aorta *Itga8-CreER^T2^* aorta. A complete list of differentially expressed genes is provided in Supplementary Table 2.

### Recombination efficiency of *Itga8-CreER^T2^* and *Myh11-CreER^T2^*

The differential expression of CreER^T2^ protein suggested there could be differences in recombination efficiency between the two *CreER^T2^* driver strains. Several independent floxed mouse models were tested for recombination efficiency between *Myh11-CreER^T2^* and *Itga8-CreER^T2^* mice. Quantitative PCR of genomic DNA before and after tamoxifen administration showed comparable recombination of the *mT/mG* reporter in aorta of both *CreER^T2^* strains (Fig. 3a, 3b). As expected, there was high-level recombination in bladder of *Myh11-CreER^T2^* mice, but very little in the bladder of *Itga8-CreER^T2^* mice (Fig. 3a, 3b). Similar findings were observed in a tamoxifen-inducible *Myocardin* mouse (Fig. 3c, 3d). Next, the activity of each *CreER^T2^* strain was quantitated in popliteal vessels. Results showed ∼90% and ∼65% recombination efficiency in lymphatic vessels of the *mT/mG* reporter in *Myh11-CreER^T2^* and *Itga8-CreER^T2^* mice, respectively (Fig. 3e, 3f). Recombination efficiencies were somewhat lower in popliteal artery and vein, but the *Myh11-CreER^T2^* activity was higher than *Itga8-CreER^T2^* (Supplementary Fig. 8). Finally, each *CreER^T2^* strain was crossed to a floxed *Srf* mouse^38^ treated with oil or tamoxifen. Normalized SRF protein was similarly knocked down in the aorta of each *CreER^T2^* strain; however, only *Myh11-CreER^T2^* showed significant knockdown of SRF in bladder (Fig. 3g, 3h). Overall, despite much lower expression of CreER^T2^ protein in *Itga8-CreER^T2^* mice, excision of floxed sequences approximates that seen in *Myh11-CreER^T2^* mice, though there appear to be context-dependent variances in recombination efficiency with different floxed alleles.

**Fig. 3.**
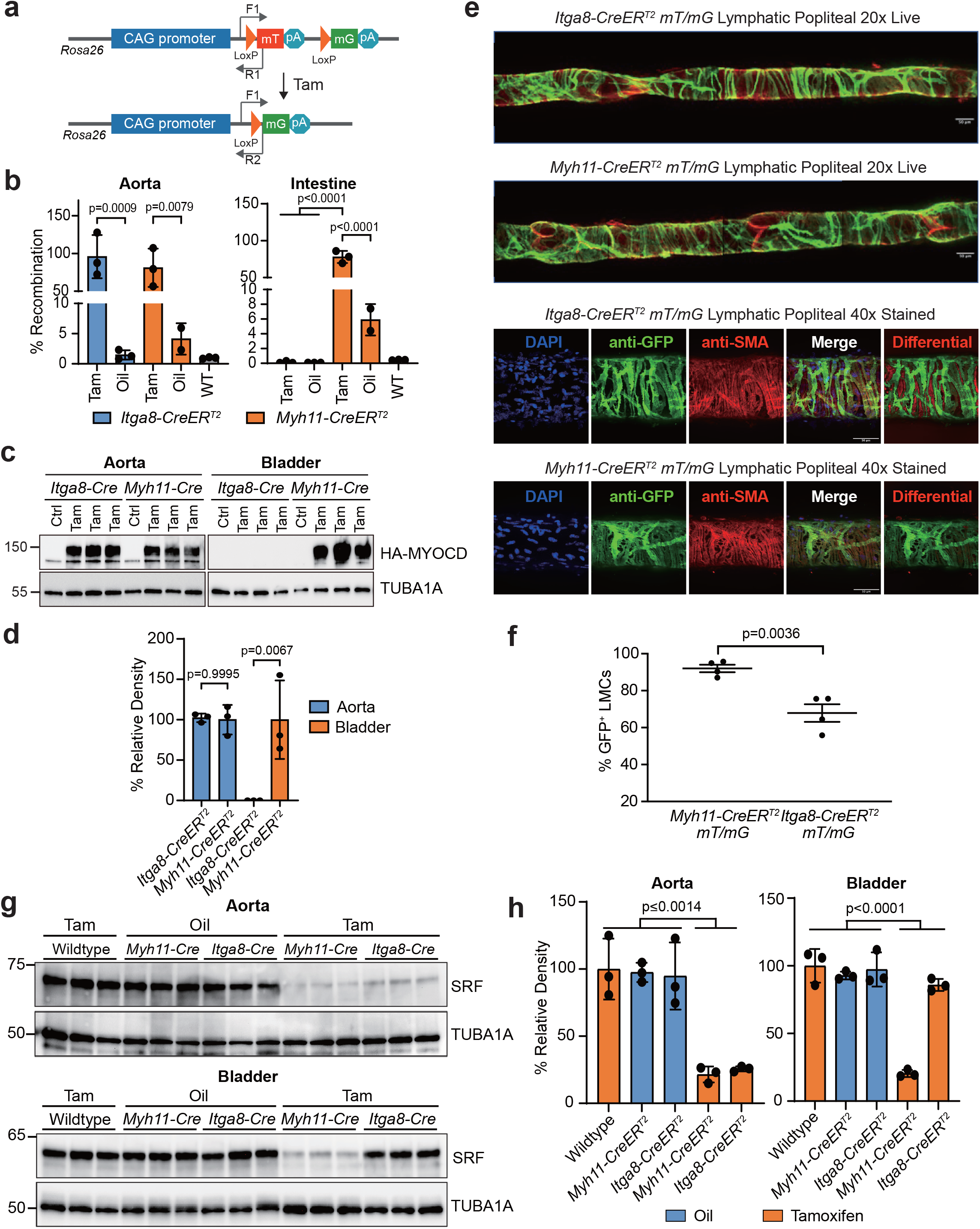
Recombination efficiency of *Itga8-CreER^T2^* versus *Myh11-CreER^T2^*. **a,** Schematic of *mT/mG* reporter at the *Rosa26* locus and position of primers (arrows) for measuring recombination. **b**, qPCR of genomic DNA following Oil or Tamoxifen (Tam) administration. **c**, Similar study as above only an inducible *Myocd* transgenic mouse was bred to each *CreER^T2^* driver for oil (Ctrl) or Tam administration and Western blotting for the presence of the HA-tagged MYOCD protein. **d**, Quantitative data of Western blots in panel **c**; percent relative density to the TUBA1A loading control; n=3 independent mice per condition. **e**, Snapshot of live images of cultured popliteal lymphatic vessels from each *CreER^T2^* and **f**, quantitation of GFP+ fluorescence in lymphatic muscle cells (LMCs); n=4 independent mice per genotype. **g**, Western blot of SRF protein in aorta and bladders of each indicated genotype with or without Tam administration. **h**, Quantitative data of panel **g**; percent relative density to the TUBA1A loading control; n = 3 independent mice per condition.

### *Itga8-CreER^T2^* circumvents a previously lethal visceral myopathy upon knockdown of *Srf*

*Srf* loss-of-function studies using an *Sm22-CreER^T2^* driver mouse demonstrated a lethal gastrointestinal phenotype.^12, 13^ Consistent with these findings, homozygous floxed *Srf* mice carrying *Myh11-CreER^T2^* displayed uniform lethality within 3-4 weeks of tamoxifen administration. Mutant mice showed a pale, distended intestine and a pronounced reduction in SRF expression within the muscular layer of the small intestine (Supplementary Fig. 9). In contrast, floxed *Srf* mice carrying the *Itga8-CreER^T2^* driver were viable with no intestinal phenotype up to 8 weeks post-tamoxifen administration (Supplementary Fig. 9 and below).

Confocal immunofluorescence microscopy demonstrated a clear reduction of SRF and ACTA2 in carotid artery of *Itga8-CreER^T2^*-mediated *Srf* knockout mice, but not in intestinal SMCs (Supplementary Fig. 10). Quantitative measures of the change in SRF protein on sections of carotid artery and intestine were consistent with confocal images (Supplementary Fig. 10). Additional Western blotting indicated a reduction in SRF of homozygous floxed (HoF) *Srf* knockout mice, but no significant change in bladder of the same mice (Supplementary Fig. 10). Notably, there was little change in SRF expression in aorta of heterozygous floxed (HeF) *Srf* knockout mice, suggesting compensatory expression from the remaining wild type allele (Supplementary Fig. 10). Because of the activity of *Itga8-CreER^T2^* in the glomerulus (Fig. 1), we considered the possibility of a kidney phenotype with conditional inactivation of *Srf*. Homozygous and heterozygous conditional *Srf* knockout mice displayed similar body weights and a normal histological appearance of kidneys, with no evidence of proteinuria (Supplementary Fig. 11). Taken together, these results establish the utility of the *Itga8-CreER^T2^* driver in evading an otherwise lethal, visceral myopathy upon inducible inactivation of *Srf* in *Myh11-CreER^T2^* mice.

### *Itga8-CreER^T2^*-mediated *Srf* knockout causes defective vascular contractile activity

To further demonstrate the utility of *Itga8-CreER^T2^*, blood pressure and vascular contractile competence were assessed in male and female mice following selective loss of *Srf*. Telemetry measures in male mice demonstrated no change in baseline blood pressure. However, following angiotensin II infusion, significant increases in systolic, diastolic, and mean arterial pressures were observed in control mice, but not in conditional *Srf* knockout mice (Fig. 4). Interestingly, age-matched female mice failed to respond to angiotensin II and showed no changes in blood pressure, regardless of *Srf* status (Fig. 5). On the other hand, the aorta of male and female mice displayed attenuated contraction in response to KCl (Fig. 6a) and phenylephrine (Fig. 6b, 6c). Immunofluorescence confocal microscopy revealed a clear decrease in SRF expression within SMCs of the aorta (Fig. 6d). Finally, bulk RNA-seq studies of the aorta from conditional *Srf* knockout mice yielded significant reduction in expression of several SMC contractile genes (Fig. 7a, 7b). Analysis of the top 250 down-regulated genes disclosed the SRF binding CArG box as the most significantly enriched transcription factor binding site (Fig. 7c). Further assessment of the down-regulated gene set revealed molecular functions related to the cytoskeleton and contractile machinery (Fig. 7d). Interestingly, the upregulated gene set following conditional *Srf* knockout was most related to metabolism (Fig. 7e). Collectively, these results demonstrate an unambiguous, viable mouse model of VSMC contractile incompetence following *Itga8-CreER^T2^*-mediated inactivation of *Srf*.

**Fig. 4.**
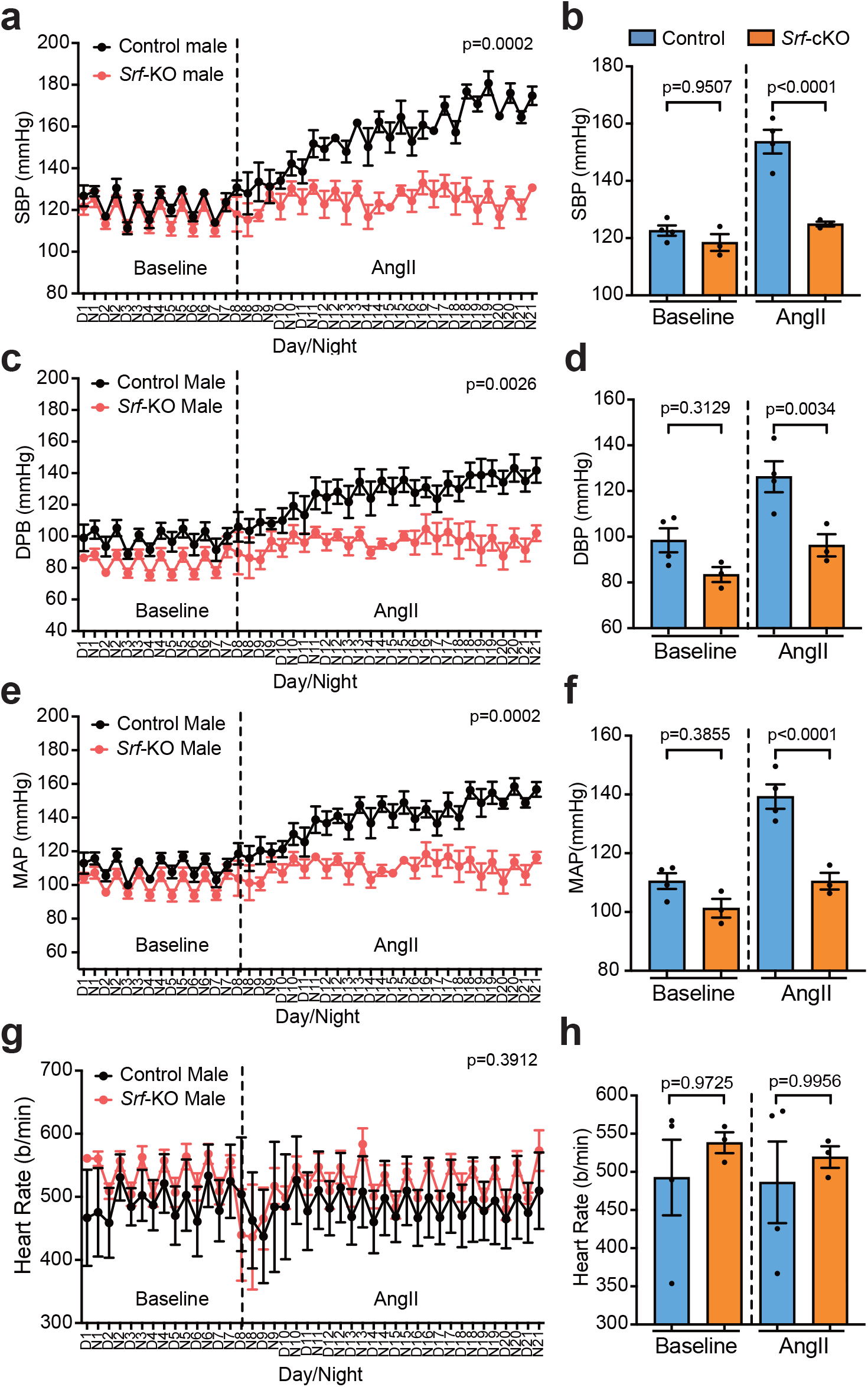
Chronic blood pressure measurements in *Itga8-CreER^T2^* mediated *Srf* knockout male mice. Effects of *Srf* conditional knockout (*Srf* cKO) on systolic blood pressure (**a, b**), diastolic blood pressure (**c, d**), mean arterial pressure (**e, f**) and heart rate (**g, h**) in male mice as measured by telemetry over 3 weeks. Control mice here and in Figures 5 and 6 are oil-treated, homozygous floxed *Srf* mice carrying *Itga8-CreER^T2^*. n = 3 or 4 mice per condition.

**Fig. 5.**
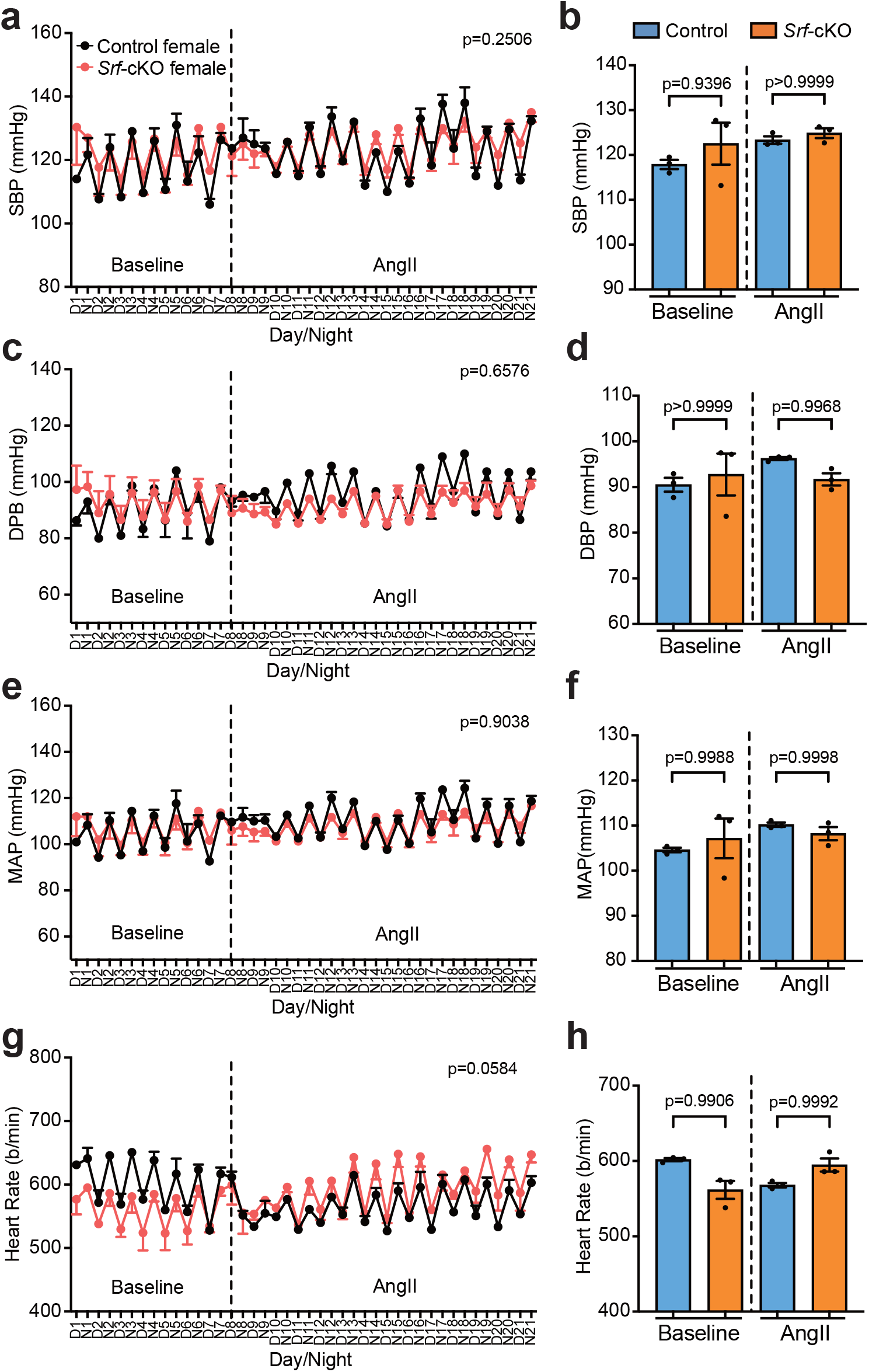
Chronic blood pressure measurements in *Itga8-CreER^T2^* mediated *Srf* knockout female mice. Effects of *Srf* conditional knockout (*Srf* cKO) on systolic blood pressure (**a, b**), diastolic blood pressure (**c, d**), mean arterial pressure (**e, f**) and heart rate (**g, h**) in female mice as measured by telemetry over 3 weeks. n = 3 mice per condition.

**Fig. 6.**
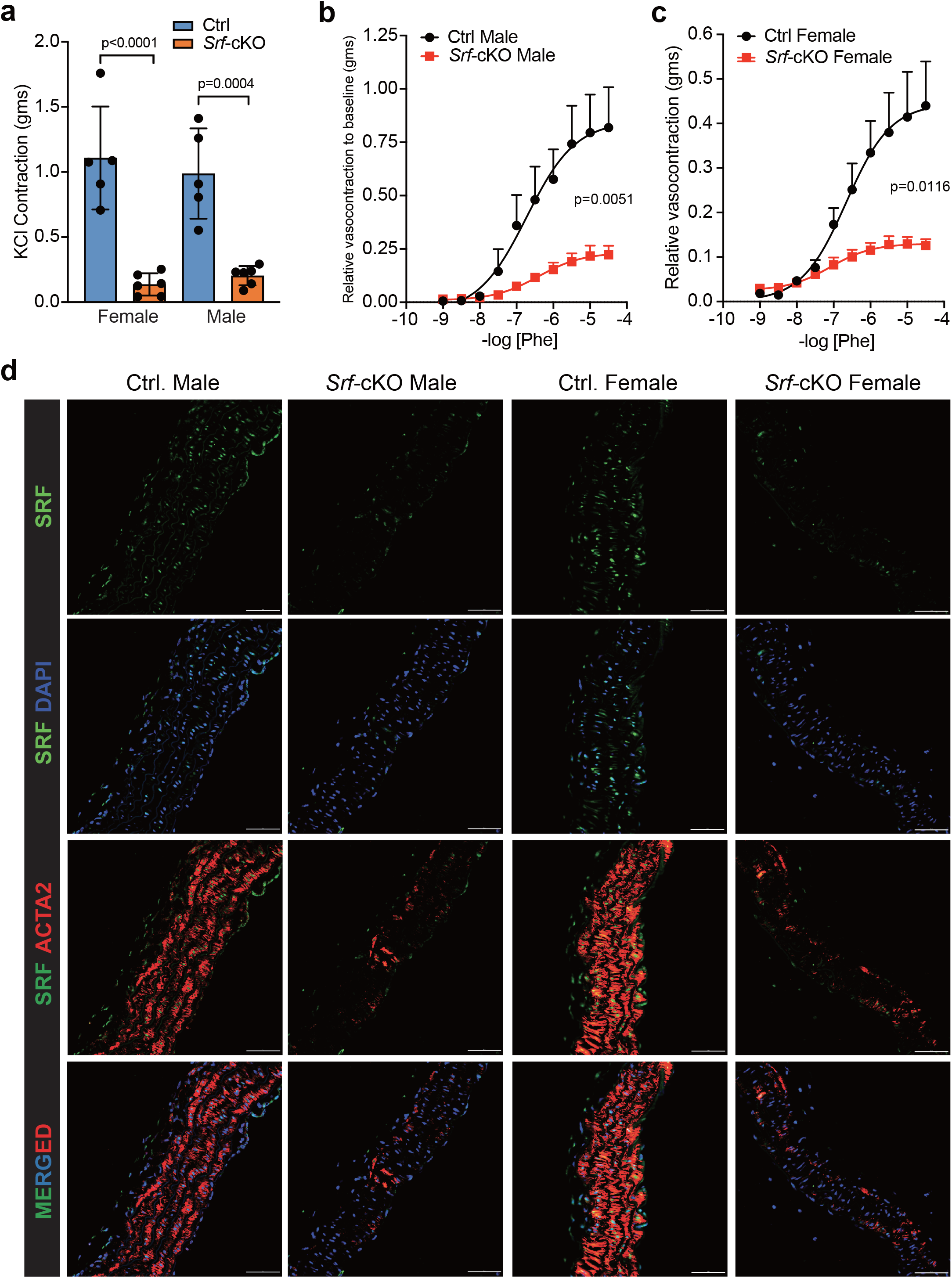
Vascular contractile competence in *Itga8-CreER^T2^* mediated *Srf* knockout mice. **a**, KCl-stimulated contractile activity in female (n=5) and male (n=6) aorta under control or cKO condition. Male (**b**) and female (**c**) aortas treated with varying doses of Phenylephrine [Phe]. **d**, Confocal immunofluorescence microscopy of indicated proteins in control and *Srf* cKO male and female aorta. The scale bars are 50 μm.

**Fig. 7.**
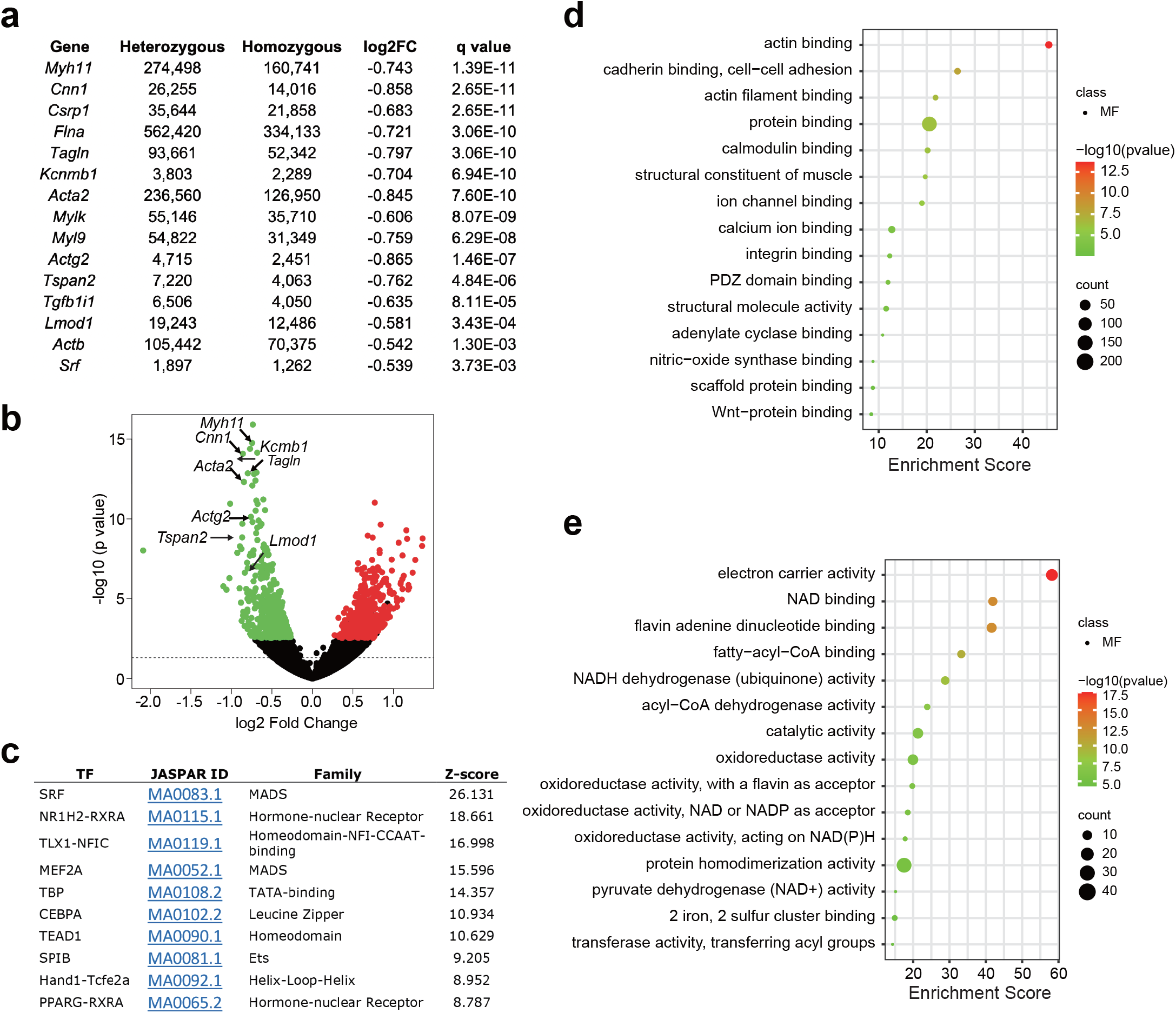
Bulk RNA-seq of aortas from *Itga8-CreER^T2^* mediated *Srf* knockout mice. **a**, Down-regulated SMC contractile genes in aortas with conditional *Srf* knockout. **b**, Volcano plot of differentially expressed genes with conditional *Srf* knockout. **c**, oPOSSUM 3.0 analysis of over-represented transcription factor binding sites in promoter/intron of top 250 down-regulated genes with conditional knockout of *Srf*. Enrichment scores for gene ontology terms (Molecular Function) related to downregulated (d) or upregulated (e) genes with conditional knockout of *Srf*.

## Discussion

All currently available SMC *Cre* mice, including the popular *Myh11-CreER^T2^* mouse,^39^ direct excision of floxed DNA in both vascular and visceral SMC lineages. Consequently, the inactivation of a critical gene in, for example, SMCs of the colon may result in a lethal phenotype and the inability to accurately interrogate gene function in VSMCs. Here, we show preferential activity of *Itga8-CreER^T2^* in SMCs of the vascular and lymphatic system with comparatively less activity in visceral SMCs of the gastrointestinal (*e.g.*, stomach and intestine) and urogenital (*e.g.*, bladder and uterus) systems. Despite low level expression of CreERT2 in *Itga8-Cre*ER*^T2^* mice, recombination of several floxed sequences occurs with no evidence of leaky activity. In contrast, *Myh11-CreER^T2^* mouse tissues show much higher levels of CreERT2 expression due to two copies of the transgene, improved codon usage for Cre and, perhaps, higher promoter activity, all of which likely promote the observed leaky activity of this SMC *Cre* driver mouse.

We also demonstrate an altered transcriptome in *Myh11-CreER^T2^* mice, including elevated transcripts of two X chromosome-linked genes that we demonstrate with long read sequencing are contiguous with the BAC-containing *Myh11-CreER^T2^* transgenes (see below). Despite several distinctions between *Myh11-CreER^T2^* and *Itga8-CreER^T2^* (Table), the *Myh11-CreER^T2^* driver has been used successfully to inactivate well over 100 floxed alleles in SMCs (Supplementary Table 1).

However, a clear advantage of *Itga8-CreER^T2^* over *Myh11-CreER^T2^* is demonstrated here for the conditional inactivation of *Srf* in adult mice. The latter *Cre* driver resulted in a lethal intestinal phenotype within 3-4 weeks of tamoxifen administration. These findings are consistent with previous reports showing comparable intestinal pathology and early demise using *Tagln-CreER^T2^* ^12, 13^ and the aforementioned *Myh11-CreER^T2^*.^16^ On the other hand, mice with the same homozygous floxed *Srf* and a similar schedule of tamoxifen administration to induce *Itga8-CreER^T2^*-mediated recombination, were viable for at least 8 weeks post-tamoxifen with no evidence of intestinal pathology. The simplest explanation for the absence of any overt intestinal phenotype is the much lower activity of *Itga8-CreER^T2^* in intestinal SMCs. This inference is based on results of the *mT/mG* reporter studies and quantitative immunofluorescence microscopy experiments showing comparable levels of SRF protein in intestinal SMCs of mice with one or both alleles of *Srf* inactivated.

The *Itga8-CreER^T2^* mouse opens the door to new experiments in VSMCs that were previously problematic because of confounding effects of gene loss in visceral SMC tissues such as intestine and bladder. In this context, although short-term (14 day) studies of conditional loss of *Srf* in VSMCs with *Sm22-CreER^T2^* have been reported, yielding insight into mechanosensitivity and blood flow^42^ as well as arterial stiffness,^43^ phenotyping occurred at a time when there is dilation of intestine, defective peristalsis, cachexia, and even death.^12, 13^ These intestinal phenotypes would certainly elicit inflammation and the release of circulating chemokines that likely impact VSMC function. Future acute *Srf* loss-of-function studies in VSMCs can now be carried out without confounding effects of gastrointestinal pathologies.

Further, the *Itga8-CreER^T2^* mouse will be of great value in assessing loss of VSMC *Srf* (and other critical genes; see Supplementary Table 1) in more chronic vascular injury models such as aneurysmal formation, atherosclerosis, transplant arteriopathy, and hypertension.

It should be noted that heterozygous *Srf* knockout mice are not haploinsufficient and exhibit both wild expression of SRF protein as well as normal tissue structure and function. This highlights an important consideration for controlling conditional knockout studies. If loss of one allele does not result in a reduction of protein expression, as seen here with SRF, then the most stringent control would be heterozygous floxed mice carrying *Cre* with the same tamoxifen schedule used in homozygous floxed mice. However, the inclusion of *Cre* and a heterozygous floxed allele may not always be the best control. For example, the *Myh11-CreER^T2^* strain studied here shows leaky CreER^T2^ activity indicating an overestimation in the number of SMCs that underwent tamoxifen-mediated excision. In addition, the altered transcriptome seen in *Myh11-CreER^T2^* mice could confound the interpretation of a phenotype. In both cases, a better control would be tamoxifen-treated *Myh11-CreER^T2^* mice carrying wild type alleles. We suggest careful analysis of floxed mice before embarking on conditional knockout studies to inform the best control model to use in experiments.

The molecular basis for leaky activity in the *Myh11-CreER^T2^* mouse is unknown. The higher levels of CreER^T2^ protein in *Myh11-CreER^T2^* over *Itga8-CreER^T2^* mice would seem to be a contributing factor. It is possible high levels of CreER^T2^ protein favor dissociation from the chaperone, HSP90, which anchors CreER^T2^ in the cytoplasm thereby prohibiting entry into the nucleus.^44^ In addition, the *Myh11-CreER^T2^* mouse has been bred far longer, with an unknown generation number, than *Itga8-CreER^T2^* allowing for a greater mutational load in the genome and the potential for modifiers influencing the trafficking of CreER^T2^. In this context, strict breeding practices should be followed with frequent refreshing of the wild type colony to prevent genetic drift and the emergence of a substrain of *Cre* driver. Whatever the mechanism(s) for a leaky *Myh11-CreER^T2^*, it is advised to routinely evaluate any *Cre* driver mice with a reporter for determination of accurate activity and to avoid continuous sibling mating for purposes of maintaining the line.

Another advantage of the *Itga8-CreER^T2^* mouse over the *Myh11-CreER^T2^* mouse is the ability to perform sex studies which enables compliance with the National Institutes of Health directive of addressing sex as a biological variable in research projects.^45^ Of note, the low activity of *Itga8-CreER^T2^* in uterine SMCs suggests this *CreER^T2^* driver may not be useful for studying effects of conditional SMC knockout studies in the female reproductive track. Although the *Myh11-CreER^T2^* mouse carries the *CreER^T2^* allele on the Y chromosome, thereby preventing studies in females, a prior report showed translocation of *Myh11-CreER^T2^* to the X chromosome and the generation of female mice that display *Myh11-CreER^T2^* activity.^46^ It should be pointed out, however, that the established t(Y;X) *Myh11-CreER^T2^* line was not characterized with respect to site-of-integration on the X chromosome^46^ and no study has yet to report the insertion site of *Myh11-CreER^T2^* on the Y chromosome. Here, using third generation sequencing, we physically mapped the *Myh11-CreER^T2^* transgene and surmise that it integrated initially at the terminal end of the X chromosome near the PAR, where high rates of recombination occur.^47^ The *Myh11-CreER^T2^* transgene then translocated to the Y chromosome carrying at least two X-linked genes, which we observe to be more abundantly expressed in *Myh11-CreER^T2^* mice. The insertion site on this “neo Y chromosome” has not been pinpointed as yet, but it must be close to the pseudoautosomal boundary on chromosome Y to explain the ∼3% of cases of t(Y;X) we and others have seen.^46^ There likely exists several forms of the *Myh11-CreER^T2^* driver due to stochastic recombination and probable variation in breakpoints that could lead to spurious reporting of a conditional knockout in SMCs. We suggest future generation of *Cre* (or other recombinase) mice use a more targeted approach for single copy integration, either within a gene of interest or safe harbor locus on an autosome.

Despite the advantages of the *Itga8-CreER^T2^* mouse allowing for sex studies in a more VSMC-restrictive manner and without the overestimation of recombination stemming from a leaky *CreER^T2^*, there are some limitations that need mention. First, as loss of both functional *Itga8* alleles results in postnatal lethality,^36^ it will not be useful to breed the *Itga8-CreER^T2^* line to homozygosity. Unless the site of integration is known, breeding to homozygosity is ill-advised since there is the possibility of disrupting a gene (coding or noncoding) or critical regulatory sequence.^48^ Second, as with all other SMC *Cre* drivers currently in use, *Itga8-CreER^T2^* exhibits activity in non-VSMC types, most notably glomerular cells of the kidney. Although attempts to define these cell types were inconclusive, previous work suggests they are likely to be mesangial cells.^28^ Thus, when using the *Itga8-CreER^T2^* mouse for conditional VSMC knockouts, some assessment of renal function should be done. In this report, there was no evidence of proteinuria and histology of the kidney in *Srf* mutant mice revealed normal glomerular and tubular structures. Whether a glomerular phenotype is manifest under stress conditions awaits further study. Third, since *Itga8* appears to be down-regulated under conditions of SMC de-differentiation,^30^ the *Itga8-CreER^T2^* mouse may not be optimal for studies requiring inducible activity following vascular insult. On the other hand, there is evidence for the up-regulation of ITGA8 protein late in intimal disease.^49^ Further work in various vascular disease models will resolve this question. Fourth, the full activity profile of *Itga8-CreER^T2^* during embryonic development and in other non-VSMC types or VSMC-related pericytes, where activity of *Myh11-CreER^T2^* has recently been demonstrated,^50^ is not known at this time. Continued characterization of *Itga8-CreER^T2^* under baseline and disease conditions will provide further insight into its utility in elucidating gene function in VSMCs. Finally, essentially nothing is known about the transcriptional/post-transcriptional control of *Itga8*. Transgenic mouse and in vitro luciferase assays failed to show the importance of a conserved SRF-binding site located ∼4-kilobases upstream of the start site of *Itga8* transcription.^33^ Remarkably, ectopic expression of the SRF coactivator, MYOCD, induced *Itga8* expression in an SRF-independent manner.^33^ Genome-wide MYOCD binding studies could illuminate novel enhancers that confer higher expression of *Itga8* in vascular versus visceral SMCs. Elucidating enhancers of *Itga8* might inform the molecular basis for preferential expression of this integrin gene in VSMCs and the further development and refinement of *Cre* driver mice for VSMC conditional knockout models.

In summary, the *Itga8-CreER^T2^* mouse allows for sex-based studies with preferential activity in VSMCs, an attribute that has yet to be demonstrated in any other SMC-restricted *Cre* driver. As such, the *Itga8-CreER^T2^* mouse represents a solution to the problem of competing phenotypes arising from the inactivation of critical genes in visceral SMCs.

### Methods Mouse models

The *Myh11-CreER^T2^* mouse (stock 019079)^39^ and *mT/mG* reporter (stock 007676)^51^ were obtained from the Jackson laboratory. The *Sm22-Cre* and floxed *Srf* strains have been described previously.^38^ To generate the *Itga8-CreER^T2^* allele, a DNA fragment containing *CreER^T2^*-*P2A*-*mCherry* coding sequence and SV40 polyadenylation signal (*CreER^T2^-P2A-mCherry-pA*) was synthesized by GENEWIZ. A 0.7-kb 5’-homology arm containing 5’-upstream and 5’-untranslated region sequences of exon 1 of *Itga8* and a 1.1-kb 3’-homology arm containing the coding sequence of exons 1-2 and intron 1-2 of *Itga8* were PCR-amplified from C57BL/6J genomic DNA using 5’-primer set (5’-GTACTGGTGAGGAAGAGATCCTGTTGTCT-3’ and 5’-CCCTCCTTCCCGAACGCTGTTCA-3’) and 3’ primer set (5’-CGATGTCTGCGGGAACCCACTGTC-3’ and 5’-GCTGTGGGTTTCTACTGGTCCCAAGCACCT-3’), respectively. The homology arms were cloned, along with an FRT flanked pGK-Neo, pGK-DTA, and *CreER^T2^-P2A-mCherry-pA*, into pBluescript SKII (+) to generate the *Itga8-CreER^T2^* knock-in construct (Figure I in the Data Supplement). The targeting construct was linearized with *AscI* and electroporated into C57BL/6J-129S6 hybrid G4 embryonic stem cells (ESCs) (a gift from Dr. Andras Nagy).^52^ The targeted *Itga8-CreER^T2^* ESC clones were confirmed by long and accurate PCR genotyping using external primers to identify the 1.0 kb 5’ and 1.3 kb 3’ targeted DNA fragments, and were microinjected into C57BL/6J blastocysts to generate mouse chimeras. By crossing the chimeras with flippase-expressing FLPeR mice in C57BL/6J background (Jackson Laboratory, 009086), the *Neo* cassette was removed to produce *Itga8-CreER^T2^* mouse line. The PCR primers used to identify the 5’- and 3’-homologous recombination events in the positively targeted ESCs and mice were: forward 5’- GCCTACACTTAAATTCTGTGTGACCTCAGAGTCATGT-3’) reverse (5’- CCATCAGGTTCTTGCGAACCTCATCACTCGT-3’), and forward (5’-CATGCTCCAGACTGCCTTG-3’) and reverse (5’-GGCAGGAGCTGAGGCCATAGCCTTCAC-3’), respectively. The final integrated construct was verified by Sanger sequencing both strands. Mice are currently in the N14 generation following successive back-crossing to wild type C57BL/6J mice. The inducible *Myocd* mouse (iMyocd) was generated in the same ESC as above with an HA-tagged Myocd_v1 cDNA^53^ cloned downstream of a CMV-driven stop-floxed cassette in the *Rosa26* locus.

### Tamoxifen administration

*Cre* mouse lines were crossed with *mT/mG* fluorescent reporter mice, a floxed *Srf* mouse,^38^ or an inducible *Myocd* mouse, as described. Inducible *Cre* activity was achieved by intraperitoneal injection of tamoxifen (Sigma-Aldrich, T5648) mixed and sonicated in 100% ethanol/sunflower seed oil (Sigma-Aldrich, S5007) at 1:9, v/v. The same dose of tamoxifen (40 mg/kg, ip) was delivered daily over a 5-day period, followed by a washout of tamoxifen for at least 10 days before experimental measures.

### Confocal microscopy

For frozen samples, fresh tissues were rapidly isolated and fixed overnight in 4% paraformaldehyde. Tissues were then cryoprotected over a second night with 30% sucrose followed by embedding in Optimal Cutting Temperature (OCT) compound (VWR, 102094-106). 5μm-7μm sections were cut with a cryostat (Leica Biosystems, CM 1950) and kept frozen until imaging was performed. Slides were removed from -20°C and warmed to room temperature. OCT compound was removed by washing with 1xPBS for 3 times. Auto-fluorescence was quenched with Vector^®^ TrueVIEW^®^ Autofluorescence Quenching Kit (Vector Laboratories, SP-8500-15) and the slides cover-slipped with ProLong™ Gold Antifade Mountant (Thermo, P36930) containing DAPI stain and air dried. Imaging was done on either an Olympus IX81 or Zeiss LSM900 confocal microscope with 20x, 40x oil (z-stacked), and 60x oil (z-stacked) objectives. Laser settings for confocal imaging were kept constant throughout. Final images were cropped to their published size in Adobe^®^ Photoshop^®^ with no digital enhancements, save the Testis panel corresponding to *Itga8-CreER^T2^* in supplementary Figure 6b; the latter panel was enhanced uniformly to increase brightness. For paraffin sections, tissues were isolated and immersion-fixed in methanol-H_2_O-acetic acid (60/30/10, v/v) overnight, processed in a benchtop tissue processor (Leica Biosystems, TP1020), and embedded (Sakura, Tissue Tek TEC 5) in paraffin blocks. Sections were cut with a microtome (Microm, HM 355 S) at 5μm and baked overnight at 58°C. Slides were then deparaffinized in a Leica^®^ Autostainer XL (Leica Biosystems, ST5010), stained with antibodies (Supplementary Table 3), quenched with Vector^®^ TrueVIEW^®^ Autofluorescence Quenching Kit (Vector Laboratories, SP-8500-15), and cover-slipped with ProLong™ Gold Antifade Mountant. Imaging was done on an Olympus IX81 or Zeiss LSM900 confocal microscope with 20x, 40x oil (z-stacked), and 60x oil (z-stacked) objectives. Final paraffin sectioned images were processed in Photoshop^®^ without any enhancements.

### Recombination efficiency

Several floxed mouse models carrying either *Itga8-CreER^T2^* or *Myh11-CreER^T2^* were treated with oil or tamoxifen as above. Following a standard 10-14 day washout period, tissues (aorta and bladder) were isolated for quantitative PCR of genomic DNA (*mT/mG* reporter) and Western blotting for the inducible *Myocardin* and floxed *Srf* mice, respectively. For popliteal vascular recombination efficiencies, mice were anesthetized with an injection of ketamine/xylazine (25/2.5 mg/ml, i.p.) and their legs shaved and cleaned. The saphenous artery and vein and the popliteal lymphatic vessel were excised from the tissue and kept in Krebs buffer with 0.5% BSA at 4°C until imaged. Each vessel was then cannulated and pressurized in an imaging chamber filled with Krebs buffer for further removal of adipose and connective tissue. Vessels were then equilibrated in calcium free solution for 15 minutes to prevent movement during live imaging. At least 2 separate fields of view of each vessel were acquired with a Yokagawa CSU-X Spinning Disc Confocal Microscope on an inverted Olympus IX81 with a Hamamatsu Flash4 camera using both 20x and 40x objectives. Each live vessel acquisition was a Z-stack from the bottom edge of the vessel to the midpoint for both the GFP (488nm) and tdTomato wavelengths (561nm).

Following the live image acquisition, the lymphatic popliteal vessels were fixed at room temp with ice cold 4% PFA for 15 minutes. The vessels were then washed with PBS and stored at 4C for subsequent immunostaining. The fixed vessels were permeabilized with PBST (0.3% triton) for 1 hour and then blocked for 3 hours with BlockAid (Thermofisher). Vessels were stained overnight with anti-GFP at 1:200 (rabbit, ThermoFisher A-11122) and anti-smooth muscle actin at 1:500 (mouse, Sigma A5228) in Blockaid buffer. The vessels were then washed with PBS and stained with Donkey anti-rabbit AF488 (A-21206, Thermofisher) and Donkey anti-mouse AF647 at 1:200 (A-31571, Thermofisher) for 1 hour at 4C. Vessels were washed in PBS and then recannulated and imaged as described above.

To assess recombination efficiency in the smooth muscle layer of the saphenous vein and artery, the number of circumferential vascular muscle cells expressing either GFP^+^ or tdTomato^+^ were tabulated from Z-stacks from 40x acquisitions. To assess recombination efficiency in the smooth muscle layer of popliteal lymphatic vessels the number of GFP^+^SMA^+^ and GFP^-^SMA^+^ lymphatic muscle cells were counted and tabulated from Z-stacks acquired at 40X. To create the differential images used as a visual aid, the RFP Z-stack and GFP Z-stack were converted to 32 bit images, the RFP Z-stack was divided by the GFP Z-stack in FIJI and a maximum projection was made of the resulting stack.

### Western blotting

Harvested cells or pulverized frozen tissues were washed with cold 1xPBS and lysed with Cell Lysis Buffer (Cell Signaling, 9803) containing complete protease inhibitor cocktail (Sigma-Aldrich, 4693132001). Protein concentration was determined (Bio-Rad, 5000202) and samples were boiled with NuPAGE LDS buffer (Thermo, NP0008). Samples were run in 4-20% SDS-PAGE gel (Bio-Rad, 4561094) in 1x Tris-Glycine running buffer (Bio-Rad, A0028). Proteins were transferred to PVDF membrane using Trans-blot Turbo transfer system and compatible transfer kit (Bio-Rad, 1704272). Membranes were blocked in blocking buffer (Bio-Rad, 12010020) and incubated with appropriate antibodies (Supplementary Table 3). Finally, chemiluminescent substrate (Bio-Rad, 1705061) and ChemiDoc imaging system (Bio-Rad, 12003154) were used for digital imaging of Western blots. Urine from *Itga8-CreER^T2^/Srf ^fl/fl^* and *Itga8-CreER^T2^/Srf ^fl/+^* mice was collected and denatured in NuPAGE LDS buffer. Samples were loaded onto SDS-PAGE gel and run in 1x Tris-Glycine running buffer. Gels were stained with Coomassie blue and images were documented as above. Purified bovine serum albumin was run alongside the urine samples and served as a positive control for the presence of protein.

### RNA isolation and real time quantitative RT-PCR

Total RNA was extracted using QIAzol^®^ Lysis Reagent (Qiagen, 79306) followed by isopropanol precipitation and 70% ethanol washing. GlycoBlue™ Coprecipitant was applied for single aortae to facilitate the handling of small quantities of RNA. Air-dried RNA pellets were dissolved in RNase-free water. Reverse transcription was performed using Bio-Rad iScript cDNA synthesis kit (Bio-Rad, 1708891) after DNase I treatment (Thermo, AM2222). Real time quantitative PCR was performed using iTaq Universal SYBR Green Supermix (Bio-Rad, 1725121) with primers listed in Supplementary Table 3. Following delta delta Ct normalization to a house-keeping control, levels of normalized gene expression were plotted as fold-change from control samples.

### Bulk RNA-seq

*Itga8-Cre/Srf ^fl/fl^* and *Itga8-Cre/Srf ^fl/+^* mice (n=3 mice each) were administered tamoxifen at six weeks of age as above. Four weeks after the first dose of tamoxifen, aortae were collected and homogenized in RNeasy Plus Micro Kit with genomic DNA eliminator columns (Qiagen, 74034). Total RNA library preparation (TruSeq stranded) and high-throughput RNA-seq were performed on an Illumina HiSeq 4000 by Genomics Research Core of the University of Rochester. Gene Ontology analysis was done with DAVID^54^ and over-representation of DNA binding sites was determined with oPOSSUM.^55^ RNA-seq data have been submitted to the Gene Expression Omnibus (GSE138824). In a separate experiment, aortas from male wild type mice or age-matched mice with either *Itga8-CreER^T2^* or *Myh11-CreER^T2^* (n=3) were processed for total RNA isolation, library preparation, and sequencing on the Illumina HiSeq 4000 at the University of Rochester. Data were analyzed as described previously^56^ and submitted to the Gene Expression Omnibus (TBA).

### Nuclear-cytoplasmic RNA fractionation for LncRNA expression

Cultured mouse vascular smooth muscle cells (MOVAS)^57^ were fractionated using Protein and RNA isolation system kit according to manufacturer’s protocol (Thermo Fisher, AM1921). Real-time qRT-PCR was done with primers to the LncRNA upstream of *Itga8* as well as *Actb* primers, used as a cytoplasmic marker (Supplementary Table 3).

### Flow Cytometry

Whole blood samples were collected from wild-type, *mTmG*, *Myh11-CreER^T2^/mTmG*, *Sm22-Cre/mTmG,* and *Itga8-CreER^T2^/mTmG* mice and centrifuged for 5 min at 2000x g at room temperature. Buffy coats were carefully transferred to a fresh 1.5ml microfuge tube. 500ul of 1x ACK Lysing Buffer (Thermo, A1049201) was added to lyse red blood cells. Samples were then centrifuged for 5 min at 3000x g at room temperature. Supernatants were carefully aspirated and the pellets were resuspended in 1x PBS. Cell suspensions were analyzed using Accuri C6 flow cytometer (BD Biosciences) in Flow Cytometry Core of University of Rochester. Signals from FITC channel were collected and gated for GFP positive cells from each experimental condition.

### Radiotelemetry recording of blood pressure

All physiological studies used male and female mice homozygous for floxed *Srf* and heterozygous for *Itga8-CreER^T2^* (± tamoxifen). Following a 10 day washout period from last tamoxifen (or oil) injection, male and female mice (13 weeks) were implanted with indwelling telemeters (carotid catheter) (DSI^©^ Model #PA-C10, New Brighton, MN) under isoflurane anesthesia as previously described.^58^ After a 7-day recovery period, baseline blood pressure (BP) (mean arterial pressure, systolic and diastolic pressure) and heart rate measurements were consciously recorded for 7 days. Mice were then implanted with subcutaneous miniosmotic pumps (Alzet, model 1002, 14 day pump 0.25 ul/hour) containing angiotensin II (Phoenix Pharmaceuticals) diluted in isotonic saline at a dose of 490ng/kg/min as previously described.^59^ Blood pressure was recorded subsequently for 14 days.

### Vascular Reactivity

Thoracic aortas from mice utilized for telemetry blood pressure recording protocol (ANGII-infused) were excised and cleaned of excess adipose tissue. Aortas were cut in 2mm rings and mounted on pins of a DMT^®^ wire myograph (Ann Arbor, MI) as previously described.^58^ Concentration response curves to phenylephrine (1nM -30μM concentrations) as well as maximum responses to KCl (80mM) were performed and recorded with LabChart^®^ analysis software (AD Instruments^®^, Colorado Springs, CO).

### Mapping and copy number determination of *Myh11*-CreER^T2^

Long-read libraries of (i) Ultra-Long DNA Sequencing (SQK-ULK001) and (ii) CAS9 Sequencing (SQK-CS9109) kits from ONT Nanopore were prepared following manufacturer’s instructions (www.nanoporetech.com). For Ultra-Long DNA libraries, ultrahigh molecular weight genomic DNA was isolated with Circulomics kits (NB-900-601-01 and NB-900-701-01) following manufacturer’s instructions (www.circulomics.com). For CAS9 libraries, high molecular weight genomic DNA was isolated with NEB kit (T3060) following manufacturer’s instructions (www.neb.com). Five males, including a male from Jackson Laboratory (www.jax.org/), were used as input for mapping following.^60^ For CAS9 Sequencing (CRISPR-LRS), crRNAs were designed with CHOPCHOP^61^ using default parameters (https://chopchop.cbu.uib.no) (Supplementary Table 3). Long-read libraries were run on R9.4.1 flow cells on a minION Mk1B or a GridION Mk1 with fast5 to fastq read conversion using guppy (v4.2.2) on MinKNOW (v20.10.3) MinKNOW Core (v4.1.2) and fast base-calling option for the base-call model and minimum Q-score of 7 option for read filtering. Fastq files were mapped to reference sequences (improved *Cre* or segments of chromosome 16) using the Long-Read Support (beta) plugin in Qiagen CLC Genomics Workbench (www.qiagen.com) which uses open-source tool minimap2. Mapped reads (informative reads) were manually queried against NCBI nr/nt, refseq genome databases (https://blast.ncbi.nlm.nih.gov/Blast.cgi) and UCSC genome browser with BLAT tool (https://genome.ucsc.edu), where a mini-tiled map of *Myh11*-*CreER^T2^* integration locus was constructed.^60^ To determine copy number of *Myh11*-*CreER^T2^*, qPCR was used as above. Genomic DNA was diluted to 50 ng for input. Two primer sets targeting *ER^T2^* of *Myh11*-*CreER^T2^* were normalized to internal control primers (Supplementary Table 3). Real time quantitative PCR conditions were the following: step1 95 °C for 3min; step 2 95 °C for 30sec, 60 °C for 30sec, 72 °C for 30sec for 40 cycles; melt curve analysis.

### Data availability

Long-read data (informative reads) have been submitted to NCBI SRA database (www.ncbi.nlm.nih.gov/sra) under BioProject number (TBA).

### Statistical analysis

A Shapiro-Wilk test was performed to determine whether data were normally distributed. Paired *t*-test was used for comparisons between experimental and control conditions or one- and two-way ANOVA for multiple group comparisons with Tukey’s post-hoc test for individual comparisons. All data analyses were performed in GraphPad Prism 8 (GraphPad Software). Results are expressed as mean ± standard deviation unless specified otherwise. Probability values of p<0.05 were considered statistically significant.

**Table.** Summary of differences between two SMC *Cre* drivers.

**Table. Summary of differences between two SMC Cre drivers**
| ***Itga8-CreER<sup>T2</sup>*** | ***Myh11-CreER<sup>T2</sup>*** |
| --- | --- |
| Single-copy knockin on Chr 2 | Multi-copy transgene on Chr Y |
| Chromosome stability | Exhibits t(Y;X) at ~ 3% of pups |
| Male-female studies | Male only studies |
| Original Cre | Codon-optimized Cre |
| Low level Cre expression | High level Cre expression |
| More stable transcriptome | Altered transcriptome |
| Undetectable Cre leakiness | Cre leakiness |
| Active primarily in VSMCs | Active in all SMC lineages |
| Circumvents visceral myopathies | Some lethal visceral myopathies |
| Active in mesangial cells of kidney | Active in cells of thymus |
| Unknown pericyte activity | Active in some pericytes |

## Acknowledgments

We thank members of the University of Rochester Medical Center Genomics Research Core for carrying out the RNA-seq experiments.

## Author Contributions

GDW developed the figures and performed Western blotting studies in Supplementary Figs. 4e and 9a and Figs. 2b and 3g-3h, bulk RNA-seq studies in Fig. 2e and Supplementary Table 2, and assisted with the physiological measurements in Figs. 4-6; JLF performed and analyzed vascular function and blood pressure data of Figs. 4, 5 and 6a-6c and wrote the methods section related to these experiments; JD performed the studies in Fig. 3c and 3d and conducted tamoxifen injections for studies blood pressure and myography measurements; ARG provided foundational work in the beginning and helped generate Supplementary Fig. 1; PG assisted with Western blotting studies of Supplementary Fig. 4a, 4b; ACY generated the data for Supplementary Fig. 3; OJS performed all histology, immunostaining, confocal immunofluorescence and bright-field microscopy; MJD and SDZ planned and performed experiments in Fig. 3e and 3f and Supplementary Fig. 8 showing recombination efficiency of *Myh11-CreER^T2^* and *Itga8-CreER^T2^* in blood and lymphatic vasculature beds, and they contributed to the writing and editing of the methods and results; CKC, ACY, and SHG did all breeding and genotyping of mice; CKC provided images to Supplementary figures 9c and 11a; CTB and TCK helped perform the telemetry experiments of Fig. 4 and 5; AA helped with the interpretation of *Myh11-CreER^T2^* mapping and the design of Fig. 2d; WBB prepared and analyzed the Sanger sequencing of the *Cre* in *Myh11-CreER^T2^* of Supplementary Fig. 7 and the long read sequence mapping of *Myh11-CreER^T2^* in Fig. 2c and 2d; AK contributed to Western blotting studies of Supplementary Fig. 4a, 4b and Fig. 2b; XL contributed to data in Fig. 2a and provided intellectual input into study design and interpretation; LG and XX designed and generated the *Itga8-CreER^T2^* mouse and developed Supplementary Fig. 2; EBC helped with the acquisition and interpretation of the blood pressure and vascular reactivity data; QL performed the studies in Supplementary Figs. 4, 5, 10b-10g, and 11b, 11d and Fig. 7; he also provided intellectual input throughout the study and finalized the figures; JMM conceived and supervised the entire project, analyzed all data, and outlined, wrote, and edited the manuscript.

## Supplemental Figure Legends

**Supplementary Fig. 1.**
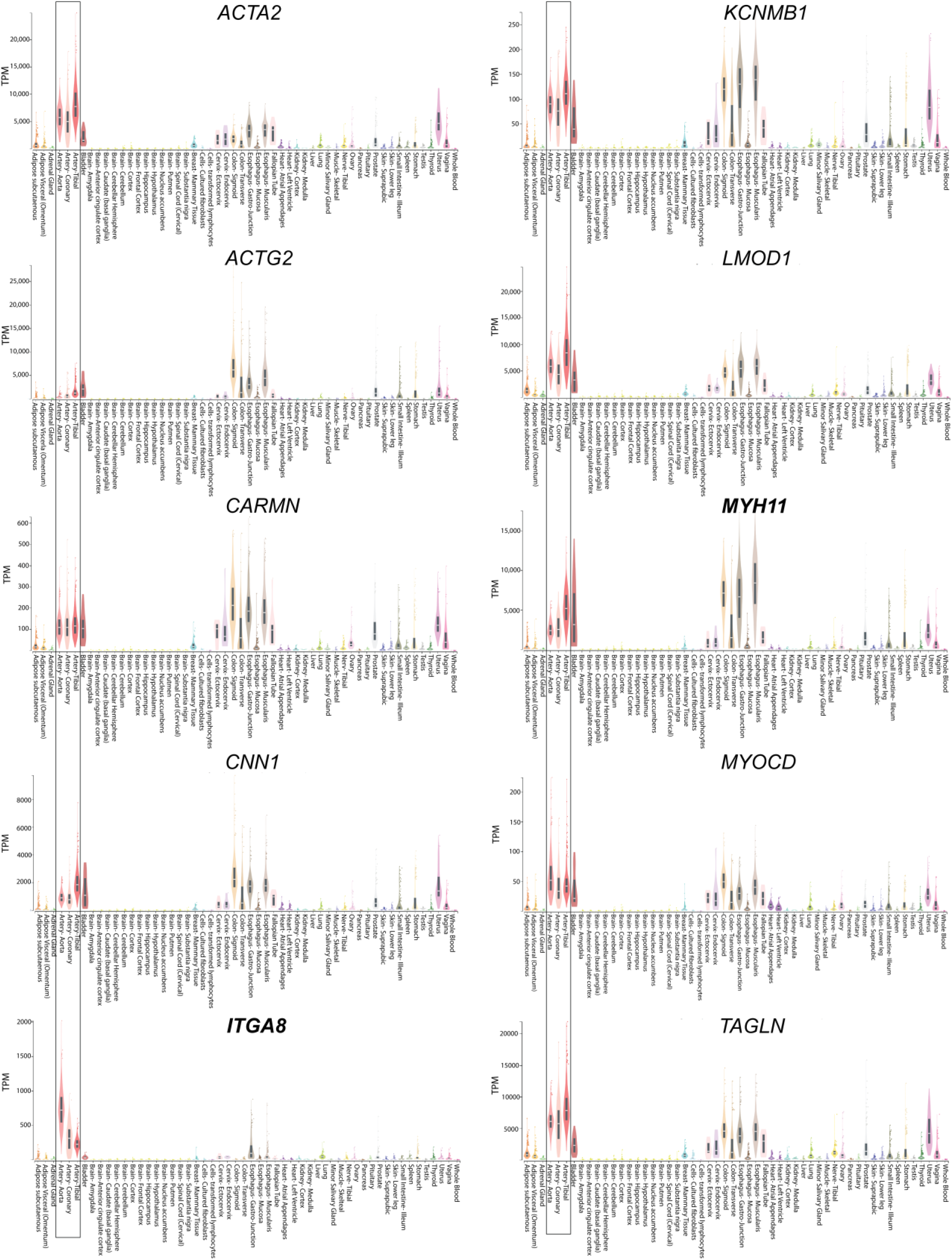
Tissue expression of indicated human genes. Screenshot of GTEx portal data of human SMC restricted genes obtained from https://www.gtexportal.org/home/gene/ITGA8. The boxed regions denote expression of each gene in the three vascular tissue types (tibial, aorta, and coronary).

**Supplementary Fig. 2.**
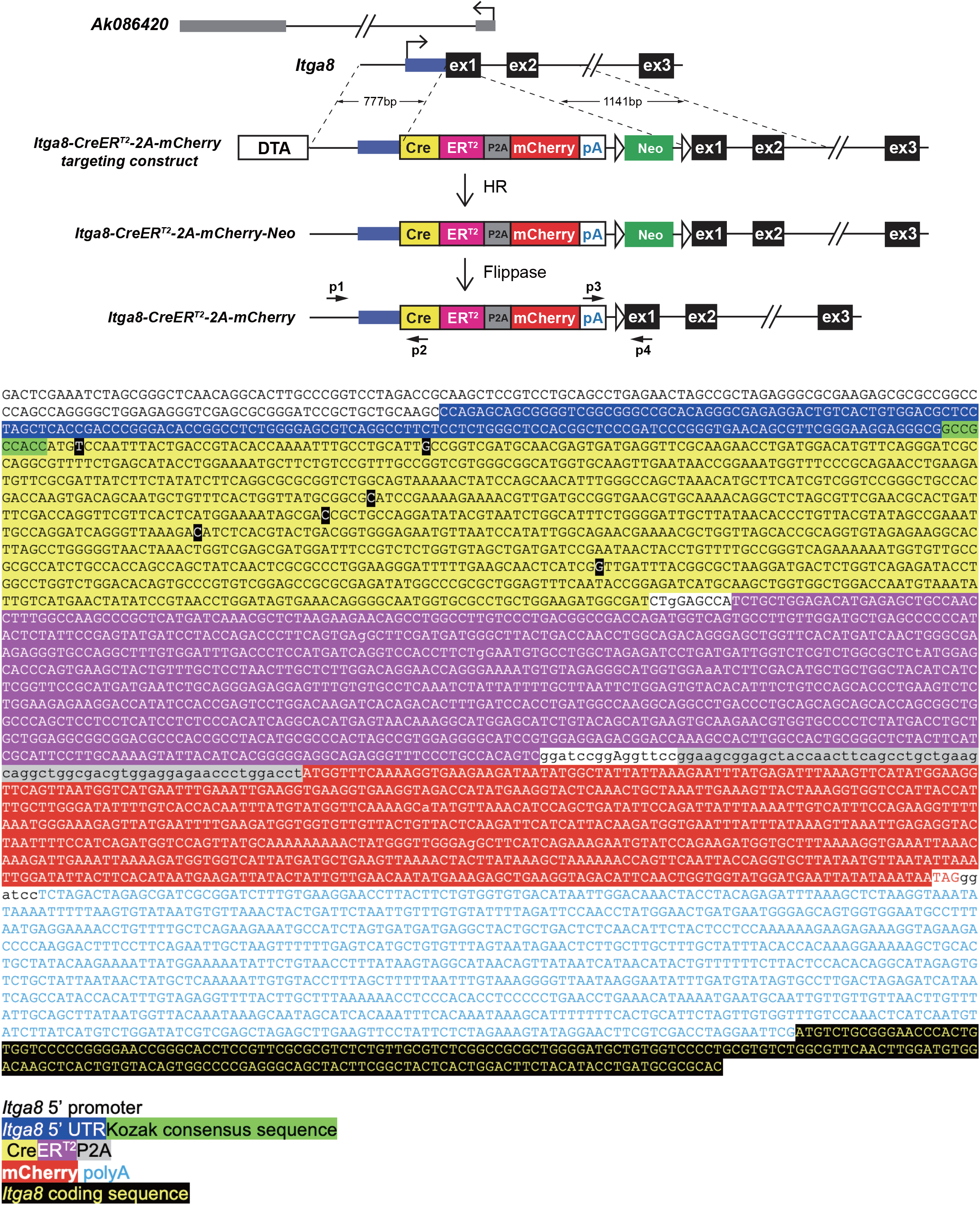
Strategy and sequence validation of *Itga8-CreER^T2^*. Top shows partial *Itga8* locus and overlapping LncRNA as well as the strategy and component parts for mouse ESC targeting. The 22 amino acid P2A “self-cleaving” cassette (GSGATNFSLLKQAGDVEENPGP) should allow for concurrent detection of the *CreER^T2^* fusion and the mCherry reporter; however, mCherry fluorescence was not pursued in this study. Small horizontal arrows denote primers for PCR genotyping of mice. Bottom shows nucleotide sequence of targeted knockin cassette following Flippase-mediated removal of the *Neo* gene; colors correspond to the schematic at top. Note Kozak consensus sequence used for optimal Cre translation. The six blackened bases in *Cre* represent codon-optimized substitutions with one non-synonymous substitution, Ala2Ser. HR, homologous recombination.

**Supplementary Fig. 3.**
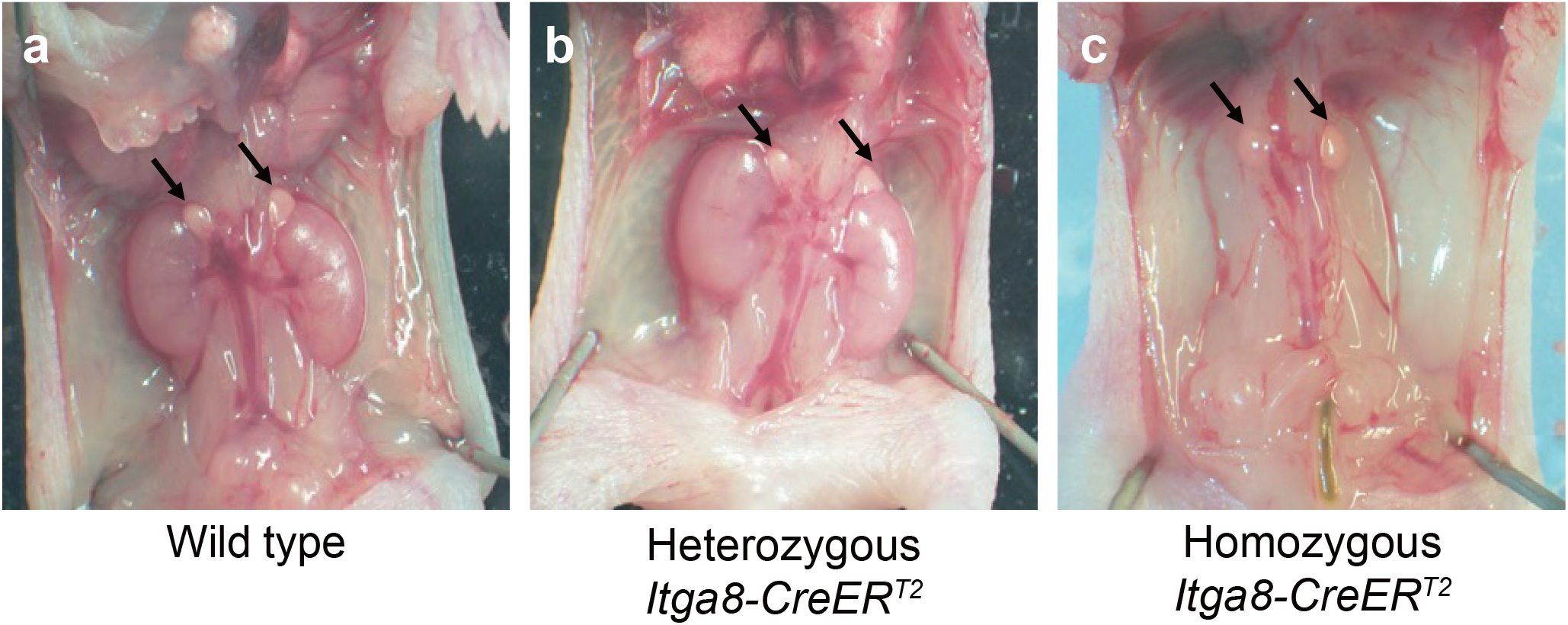
Kidney agenesis in homozygous *Itga8-CreER^T2^* mice. Arrows denote adrenal glands. Note complete loss of kidneys in the homozygous mouse.

**Supplementary Fig. 4.**
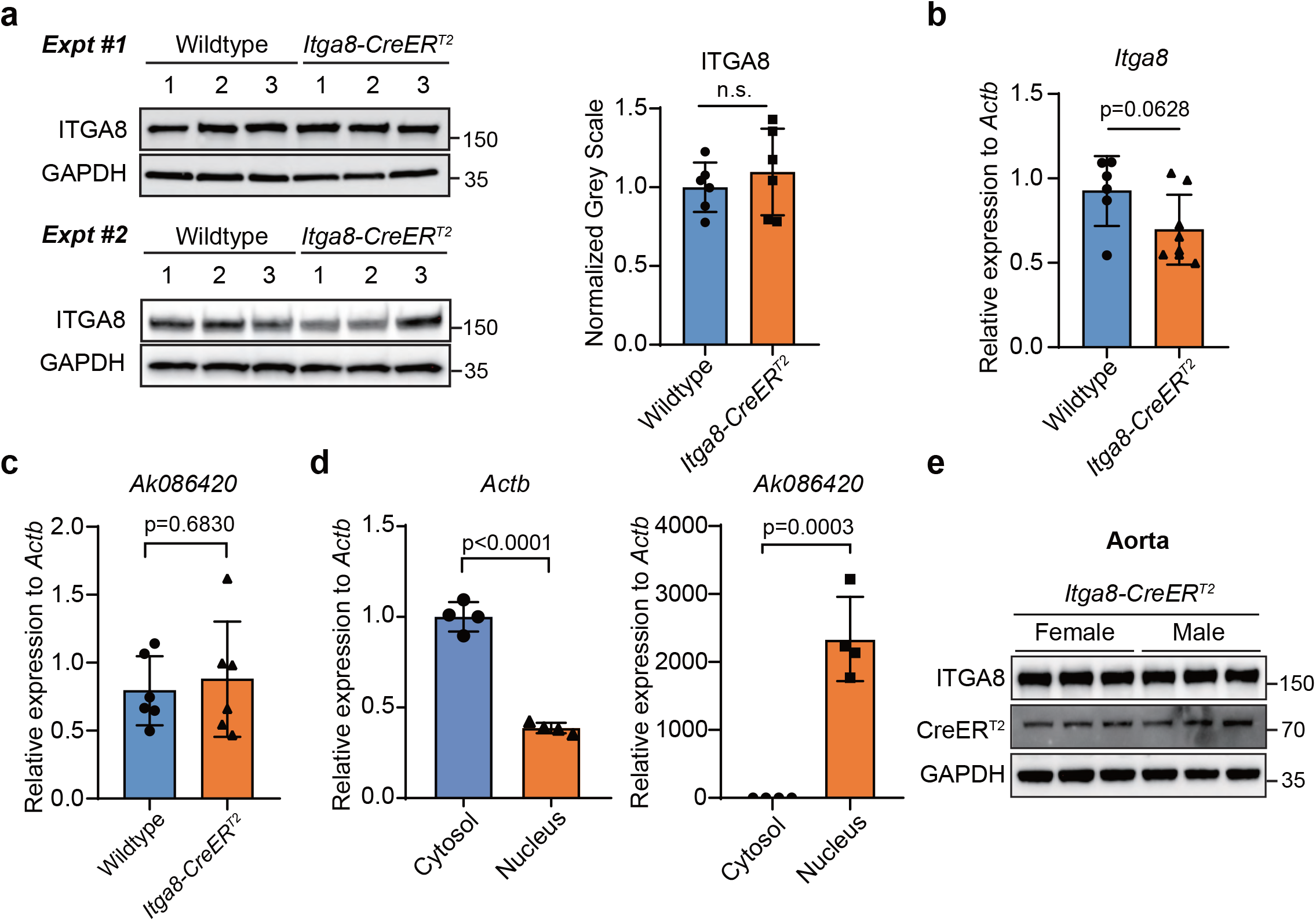
ITGA8 and LncRNA expression in heterozygous *Itga8-CreER^T2^* mice. **a**, Western blotting of ITGA8 protein in two independent experiments each in triplicate with quantitative data at right. qRT-PCR analysis of *Itga8* mRNA, **b** and the antisense *Ak086420* LncRNA, **c** in wild type (WT) versus *Itga8-CreER^T2^* heterozygous aorta. **d**, qRT-PCR of cytosolic versus nuclear gene expression in wild type aorta. n = 4 or more replicates per condition. e, Western blotting of ITGA8 and CreER^T2^ in aorta of age-matched male and female *Itga8-CreER^T2^* heterozygous mice.

**Supplementary Fig. 5.**
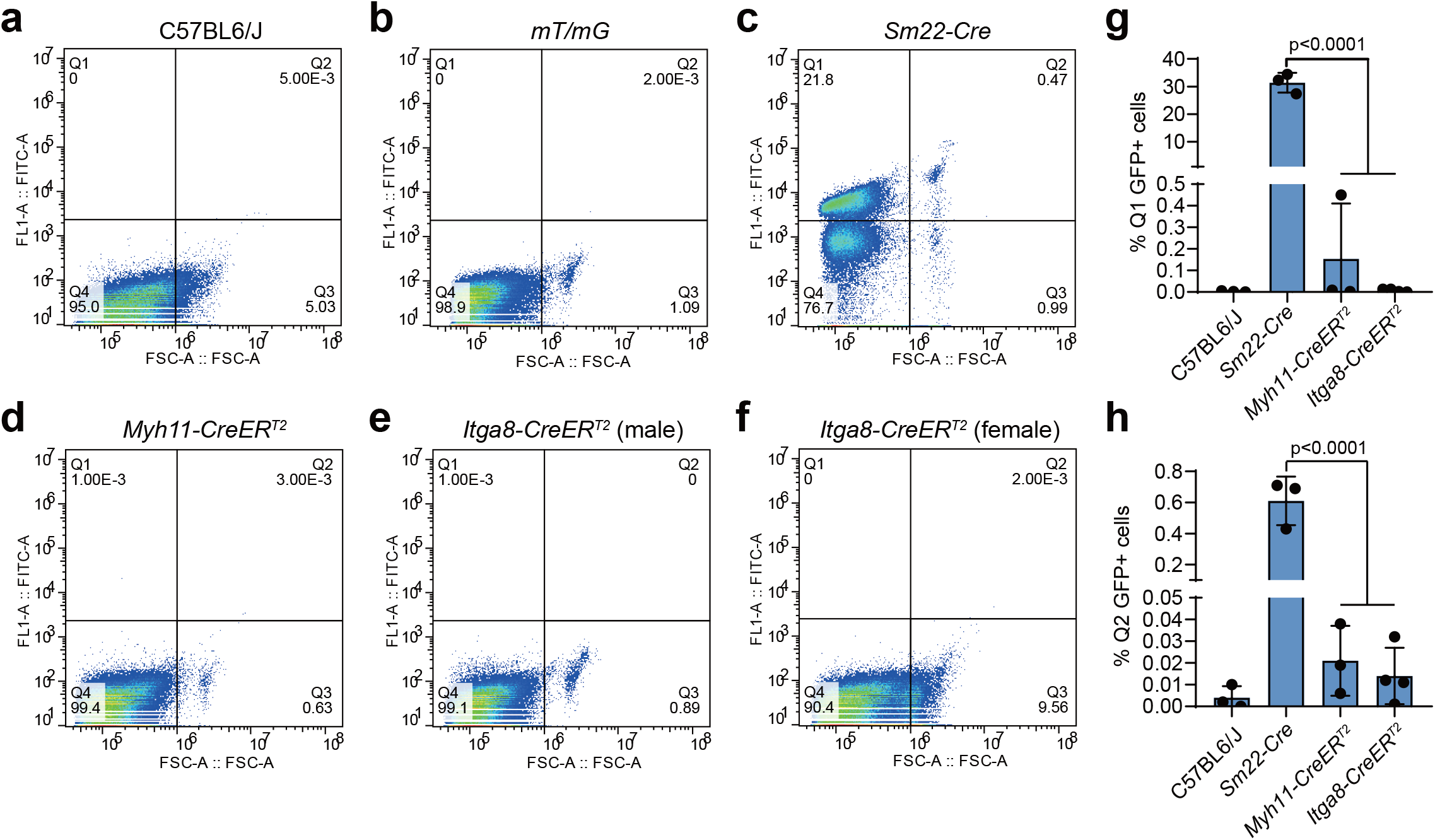
Activity of different SMC *Cre* mice in circulating cells. Flow cytometry of GFP labeled cells from **a**, wild type C57BL/6J mice; **b**, *mTmG* reporter mice; **c**, *Sm22-Cre* mice; **d**, *Myh11-CreER^T2^* mice; **e**, male *Itga8-CreER^T2^* mice; **f**, female *Itga8-CreER^T2^* mice. **g-h,** Quantitative data for GFP+ cells in upper left quadrant Q1 (**g**) and upper right quadrant Q2 (**h**) is shown for each *Cre* driver line (n=3 mice/line).

**Supplementary Fig. 6.**
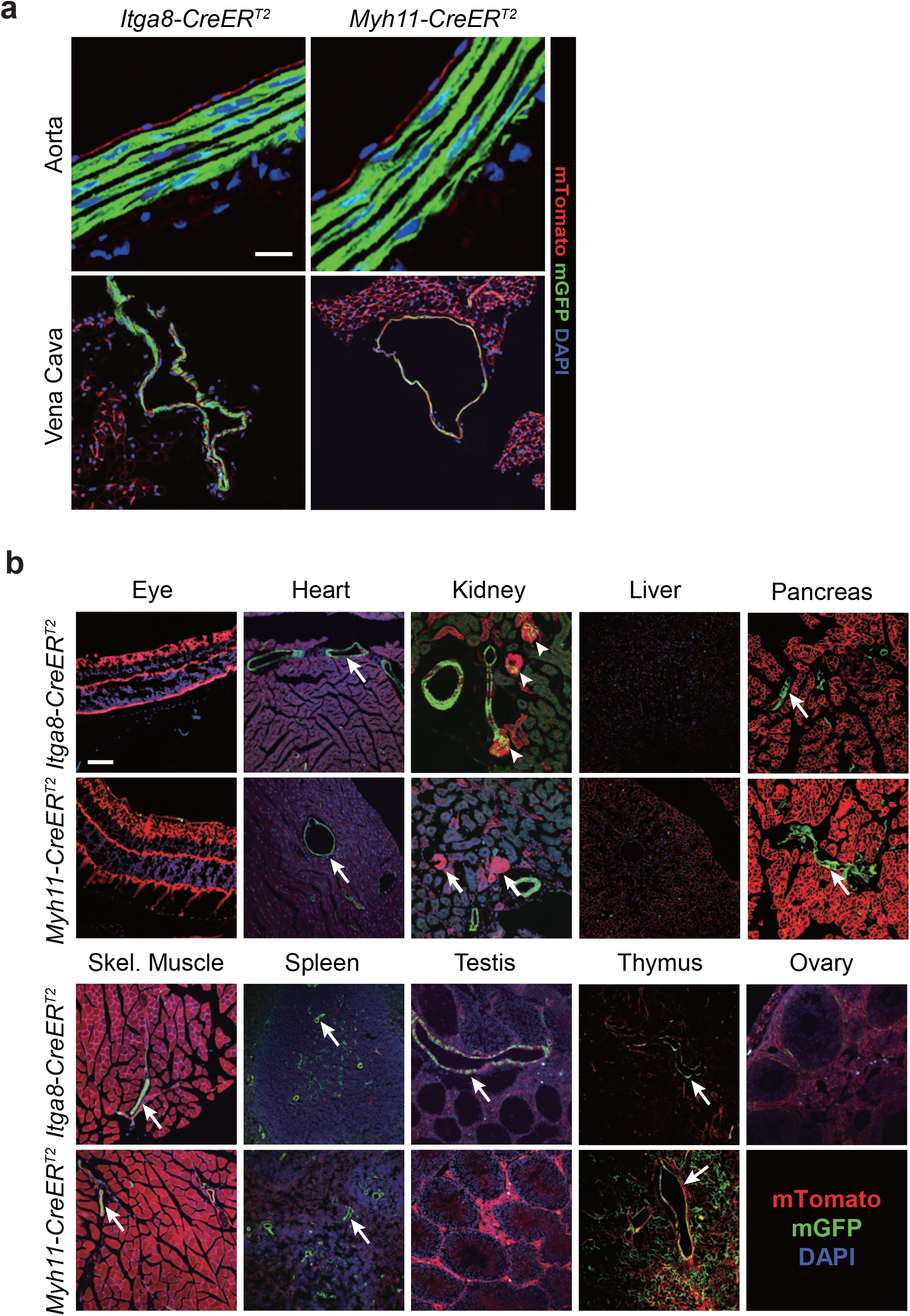
Comparative *Cre* activity in additional tissues. **a**, The GFP signal demonstrates recombination of the *mT/mG* reporter in medial SMCs of both the Aorta and Vena Cava, whereas the red stained endothelium indicates the absence of recombination. Note the absence of autofluorescence in elastic lamellae of Aorta (see methods for how this was achieved). Scale bar is 33μm for Aorta and 100μm for Vena cava. The aorta and vena cava of *Itga8-CreER^T2^* was from a female mouse. **b**, GFP signal largely restricted to vascular SMC in blood vessels (arrows) of each indicated tissue type. Arrowheads represent GFP+ cells of the glomerulus in the kidney of an *Itga8-CreER^T2^* mouse. Note diffuse GFP+ signal in thymus of *Myh11-CreER^T2^* mouse, but not *Itga8-CreER^T2^* mouse. All images were processed exactly the same except for the Testis panel under *Itga8-CreER^T2^*; here the image was enhanced in Photoshop to bring out more of the overall signal that otherwise would be too dark to visualize. Scale bar is 100μm for all images, save kidney where scale is 50μm.

**Supplementary Fig. 7.**
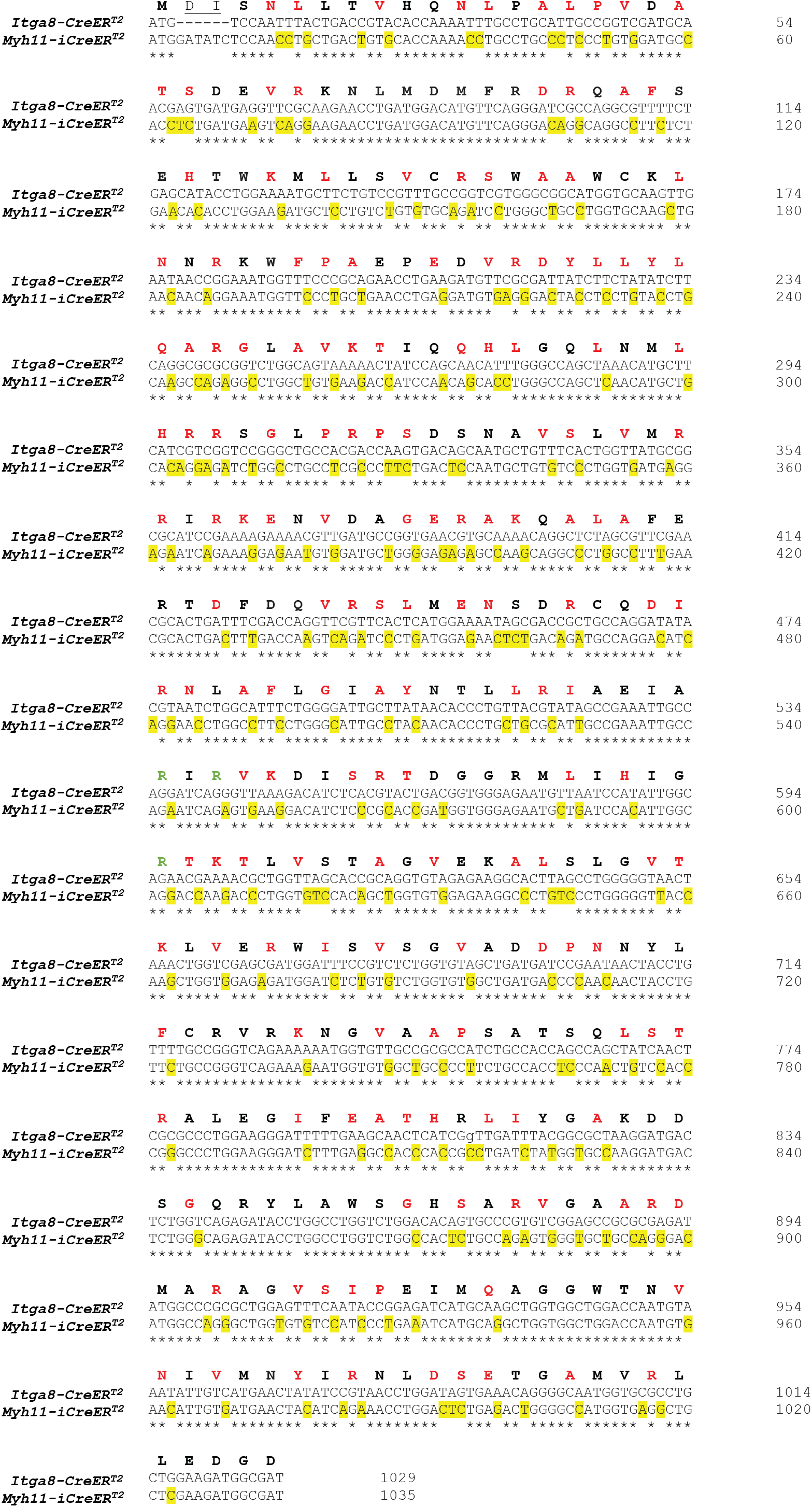
Different *Cre* sequences between *Myh11-CreER^T2^* and *Itga8-CreER^T2^* mice. Clustalw alignment of Sanger sequenced *Cre* for each driver mouse. Note *Myh11-iCreER^T2^* here (only) to indicate the improved codon usage for this *Cre* driver mouse. BLAST indicated 22% nucleotide substitutions in *Myh11-iCreER^T2^* (yellow shaded) yielding 48% of codons optimized (red letters).

**Supplementary Fig. 8.**
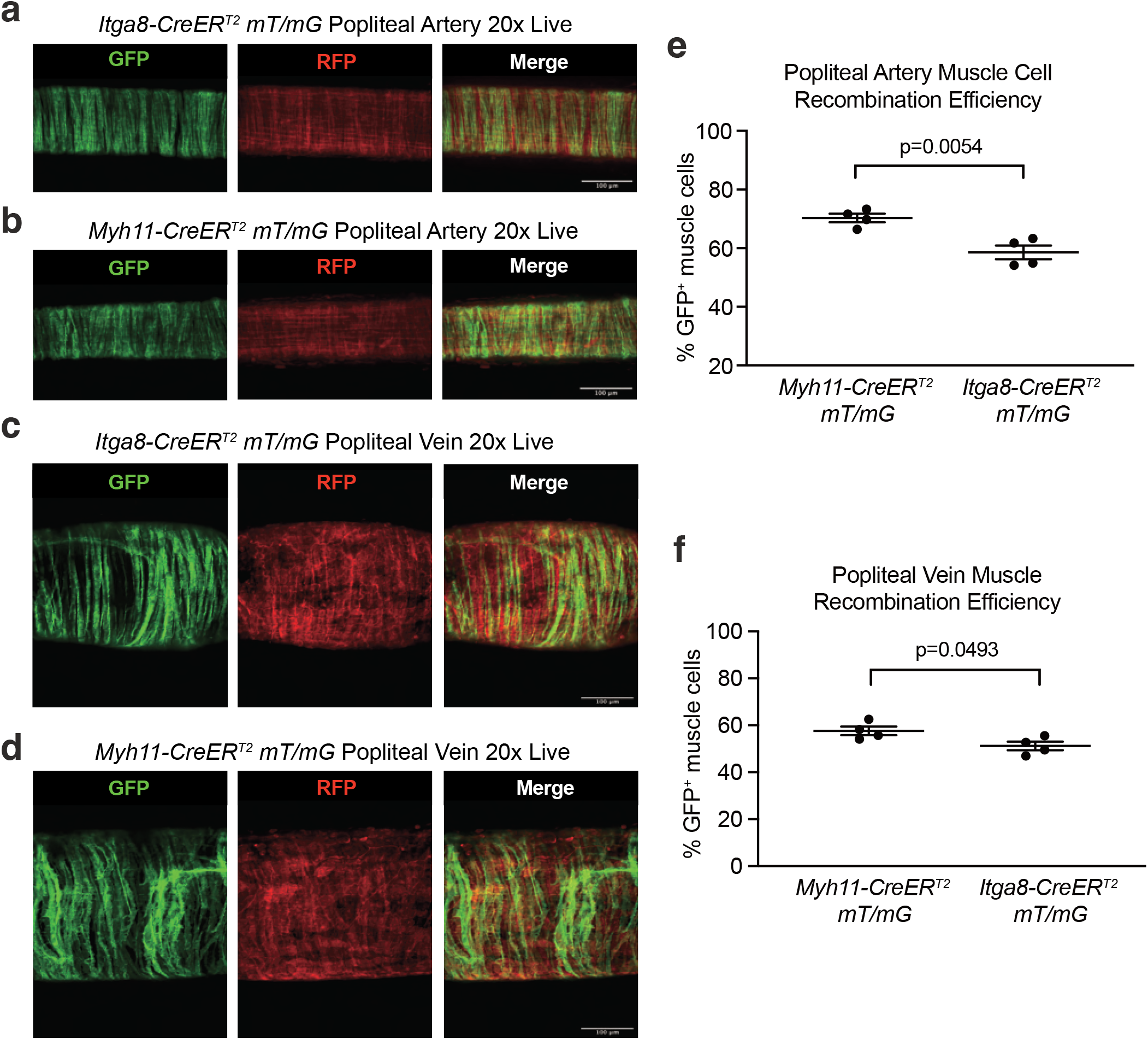
Quantitative activity of *Myh11-CreERT2* versus *Itga8-CreER^T2^* in popliteal blood vessels. Confocal imaging of artery (**a,b**) and vein (**c,d**) with quantitation of each in panels **e** and **f**, respectively. Scale bars represent 100 μm.

**Supplementary Fig. 9.**
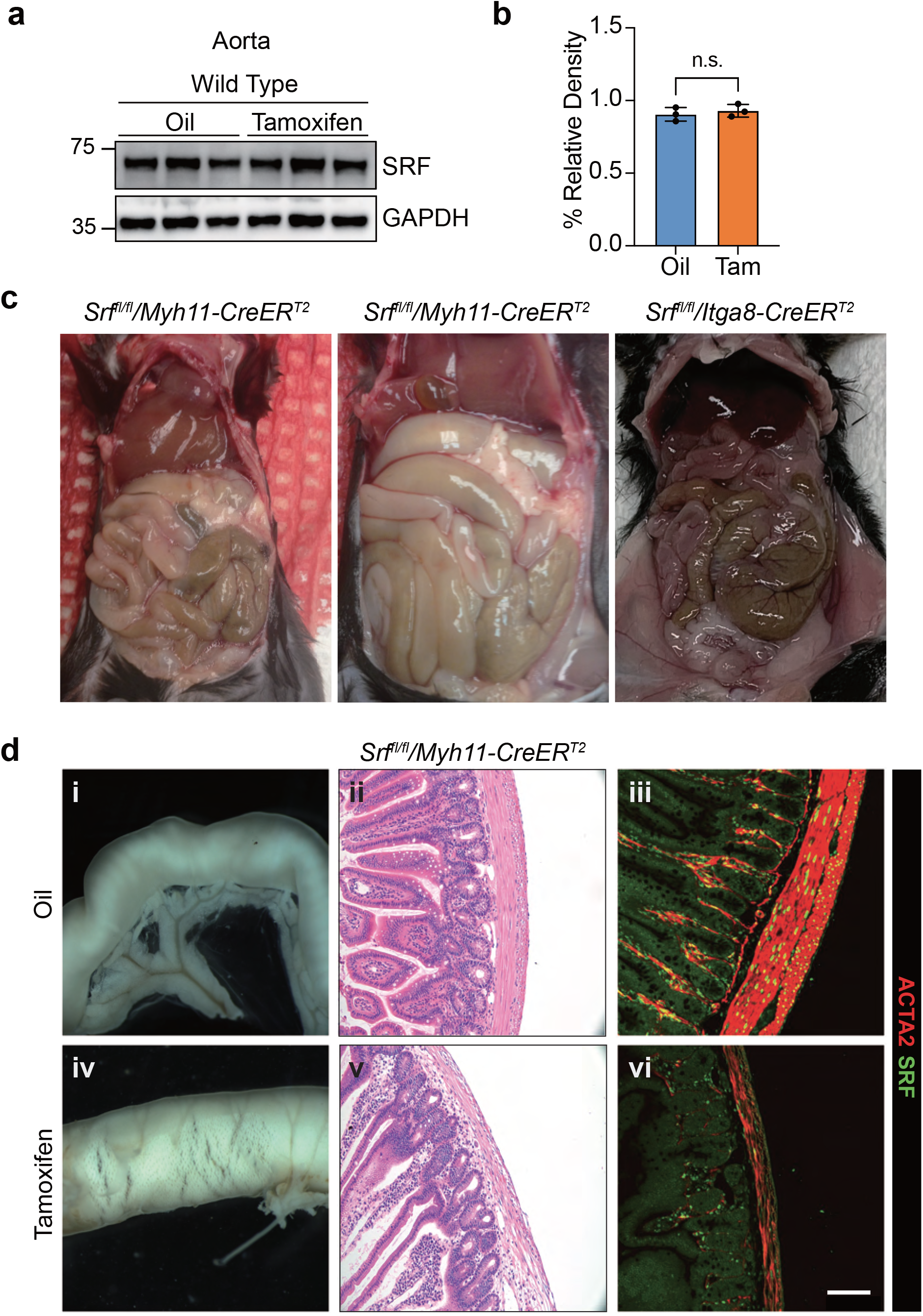
Intestinal phenotype in *Myh11-CreER^T2^* versus *Itga8-CreER^T2^* mediated knockout of serum response factor (*Srf*). **a**, Western blot showing lack of effect of tamoxifen on SRF protein levels in wild type aorta. **b**, Quantitation of panel **a**. **c**, Anatomy of abdominal cavity in the indicated genotypes of mice where *Srf* was inactivated. The two *Srf^fl/fl^*/*Myh11-CreER^T2^* images were from mice 14 days following tamoxifen administration, whereas the *Srf^fl/fl^*/*Itga8-CreER^T2^* image was from a mouse 8 weeks after tamoxifen administration. **d**, Oil (panels i-iii) and Tamoxifen (panels iv-vi) treated *Srf^fl/fl^*/*Myh11-CreER^T2^* mice with dissected gross intestine (i, iv), H&E stained intestine (ii, v), and immunostaining for SRF (green) and ACTA2 (red) in intestine (iii, vi). Scale bar is 100μm for ii, iii, v, and vi or 1mm for i and iv.

**Supplementary Fig. 10.**
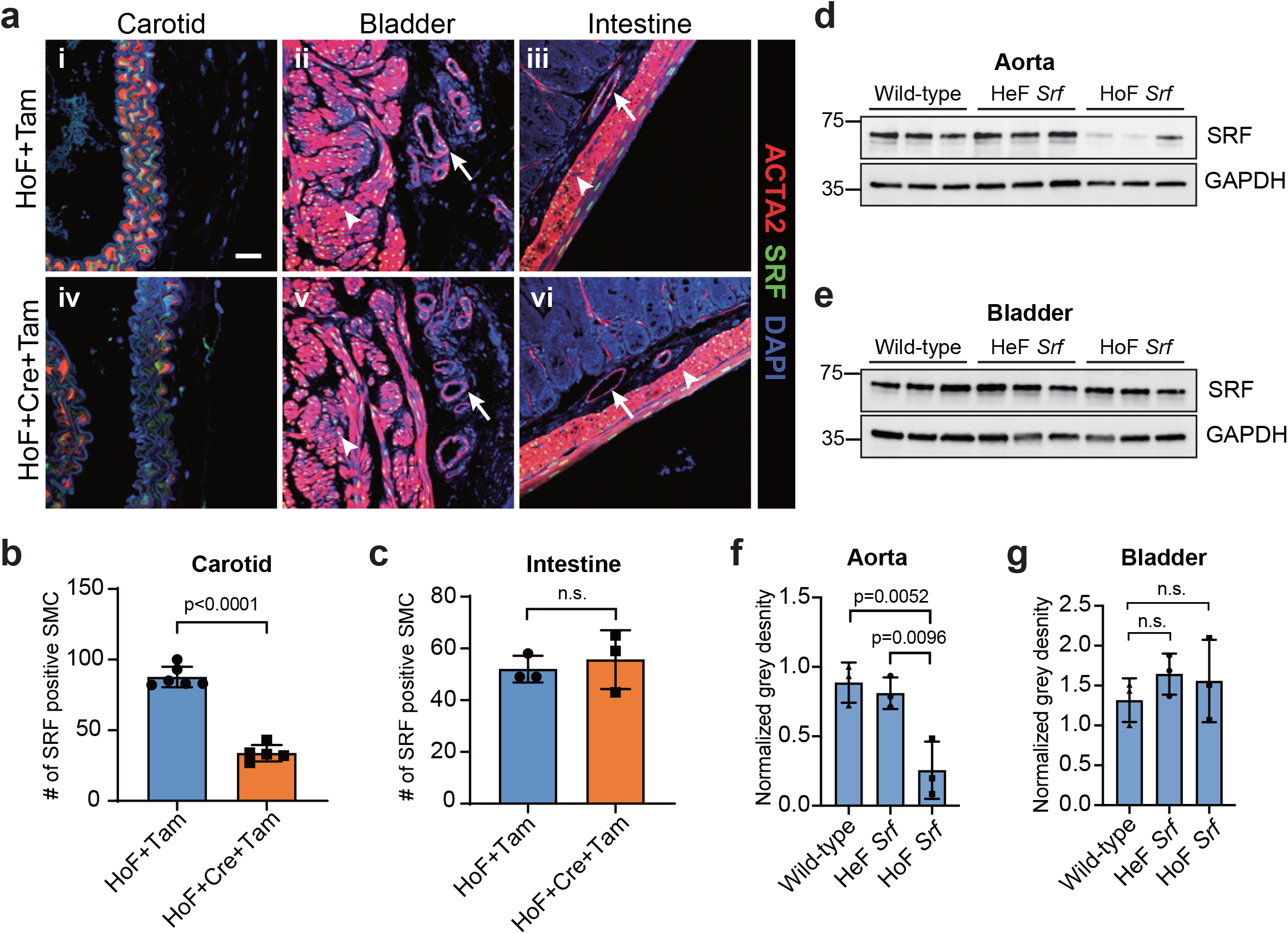
*Itga8-CreER^T2^* mediated inactivation of *Srf* in adult mouse tissues. **a**, Immunofluorescence confocal microscopy of sections of carotid artery (i, iv), bladder (ii, v), and intestine (iii, vi) from tamoxifen-treated male mice carrying homofloxed *Srf* alleles in the absence (i-iii) or presence (iv-vi) of *Itga8-CreER^T2^* (abbreviated Cre). Similar results were seen in the aorta and in vessels where one floxed *Srf* allele was excised with *Itga8-CreER^T2^* (data not shown). Sections were stained with antibodies to ACTA2 (red), SRF (green), and DAPI. Arrows and arrowheads point to blood vessels and visceral SMC, respectively. Scale bars is 20μm. SRF positive nuclei were counted in sections of tamoxifen-administered homozygous floxed *Srf* mice without *Cre* (HoF+Tam, n=6 mice) or homozygous floxed *Srf* mice with *Cre* (HoF+Cre+Tam, n=5 mice) carotid arteries (**b**) and intestine (n=3 mice each) (**c**). Western blots of SRF in aorta (**d**) and bladder (**e**) of indicated genotypes, all treated with same schedule of tamoxifen. Corresponding quantitative data is shown for mouse aorta (**f**) and bladder (**g**) n=3 mice/genotype. HeF, heterozygous floxed *Srf*; HoF, homozygous floxed *Srf*. n.s., not significant.

**Supplementary Fig. 11.**
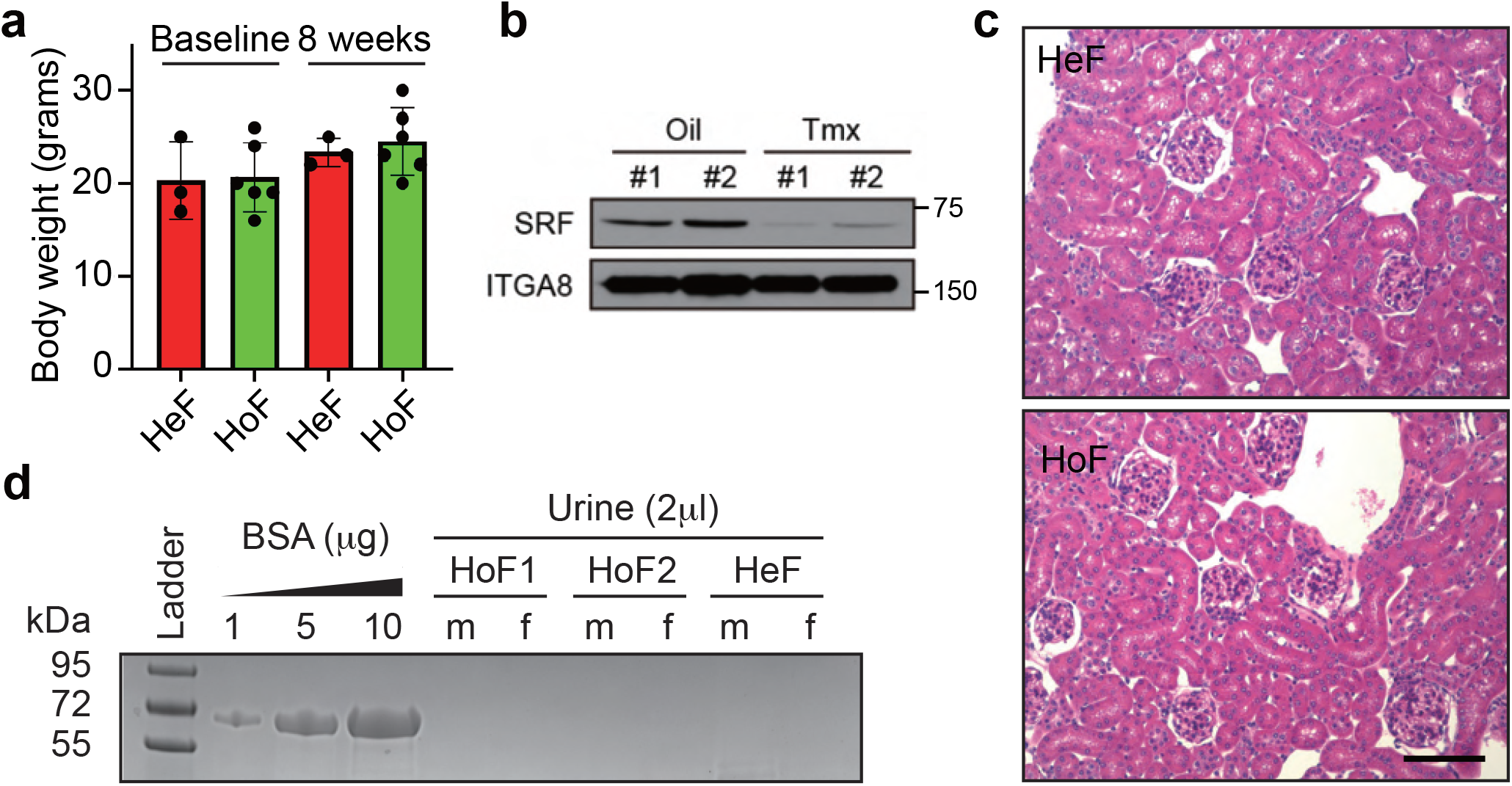
Normal body weight and kidney in tamoxifen-treated *Srf^fl/fl^/Itga8-CreER^T2^* mice. **a**, Body weights at baseline and 8 weeks post-tamoxifen treatment in heterozygous *Srf* knockout (HeF; n=3) and homozygous *Srf* knockout (HoF; n=6) mice. **b**, Western blot of ITGA8 and SRF protein in aorta of *Srf^fl/fl^/Itga8-CreER^T2^* mice treated with Oil or tamoxifen (Tmx) 6 weeks after initial dosing. **c**, H&E staining of kidney in HeF versus HoF mice 6 weeks after Tmx administration. Scale bar is 50μm. **d**, Coomassie-stained acrylamide gel of urine samples from male (m) and female (f) mice of indicated *Itga8-CreER^T2^*-mediated floxed *Srf* knockout genotypes. Bovine serum albumin (BSA) was used as a control for visualizing evidence of proteinuria.

