## Supplemental Table 3 for "An Inducible *Cre* Mouse with Preferential Activity in Vascular Smooth Muscle Evades a Previously Lethal Intestinal Phenotype"

**Supplementary Table 2. Major Resources**

**Animals (*in vivo* studies)**

| **Species** | **Vendor or Source** | **Background Strain** | **Sex** |
| --- | --- | --- | --- |
| Mice, *Mus Musculus* | Jackson Labs | C57BL/6J | M, F |
| mTomato/mGFP | Jackson Labs | C57BL/6J | M, F |
| *Sm22-Cre* | Miano Lab | C57BL/6J | M |
| *Myh11-CreER^T2^* | Jackson Labs | C57BL/6J | M |
| *Srf ^fl/fl^* | Miano Lab | C57BL/6J | M, F |
| *Itga8-CreER^T2^* | University of Rochester (URMC) | C57BL/6J | M, F |
| Inducible *MYOCD* | University of Rochester (URMC) | C57BL/6J | M |

**Animal breeding**

|  | **Species: mice** | **Vendor or Source** | **Background Strain** | **Other Information** |
| --- | --- | --- | --- | --- |
| **Parent – Male** | Wild-type mice | Jackson Labs | C57BL/6J |  |
| **Parent – Female** | Wild-type mice | Jackson Labs | C57BL/6J |  |
| **Parent – Male** | *Sm22-Cre* | Miano lab | C57BL/6J |  |
| **Parent – Female** | *mT/mG* | Jackson Labs | C57BL/6J |  |
| **Parent – Male** | *Myh11-Cre* | Jackson Labs | C57BL/6J |  |
| **Parent – Female** | *mT/mG* | Jackson Labs | C57BL/6J |  |
| **Parent – Male** | *Itga8-Cre* | URMC | C57BL/6J |  |
| **Parent – Female** | *mT/mG* | Jackson Labs | C57BL/6J |  |
| **Parent – Male** | *Itga8-Cre* | URMC | C57BL/6J |  |
| **Parent – Female** | *Srf ^fl/fl^* | Miano lab | C57BL/6J |  |
| **Parent – Male** | *mT/mG* | Jackson Labs | C57BL/6J |  |
| **Parent – Female** | *Itga8-Cre* | URMC | C57BL/6J |  |

**Primers**

| **Primer** | **Sequence** | **Usage** |
| --- | --- | --- |
| *Itga8* Fwd | TGACACCACCAACAACAGG | qRT-PCR |
| *Itga8* Rev | AGTTCTCCAGTGATACAAAGGG | qRT-PCR |
| *AK086420* Fwd | TCTTGGTTGAGCCTCATTAGTT | qRT-PCR |
| *AK086420* Rev | TCACCTCCCAGACCATTTATTC | qRT-PCR |
| *Actb* Fwd | GAGGTATCCTGACCCTGAAGTA | qRT-PCR |
| *Actb* Rev | CACACGCAGCTCATTGTAGA | qRT-PCR |
| *mT/mG* Fwd 1 | GGTTCGGCTTCTGGCGTGTGACC | Recombination efficiency qRT-PCR |
| *mT/mG* Rev 1 | GCGCATGAACTCTTTGATGAC | Recombination efficiency qRT-PCR |
| *mT/mG* Rev 2 | TCCTGTCCGTTCGCTTTG | Recombination efficiency qRT-PCR |
| iCre Fwd 1 | TGGATTTGACCCTCCATGAT | Copy number PCR 1 |
| iCre Rev 1 | GATCTCCACCATGCCCTCTA | Copy number PCR 1 |
| iCre Fwd 2 | CTTGCTCTTGGACAGGAACC | Copy number PCR 2 |
| iCre Rev 2 | TCCAGAGACTTCAGGGTGCT | Copy number PCR 2 |
| Internal Control Fwd | CACCGGCTACACCAATCAA | Copy number PCR 1 & 2 |
| Internal Control Rev | CAGAAGATGTCTCCAGGCAAG | Copy number PCR 1 & 2 |
| *Flox Srf* Fwd | TGCTTACTGGAAAGCTCATGG | Genotyping |
| *Flox Srf* Rev | TGCTGGTTTGGCATCAACT | Genotyping |
| *Itga8-Cre* Fwd | CCATGAGTGAACGAACCTGGTCG | Genotyping |
| *Itga8-Cre* Rev | GAAATCTAGCGGGCTCAACA | Genotyping |
| *mT/mG* Fwd | CTCTGCTGCCTCCTGGCTTCT | Genotyping |
| *mT/mG* Rev | CGAGGCGGATCACAAGCAATA | Genotyping |
| *Sm22-Cre* Fwd | GCATCTCCAAAGCATGCAGAG | Genotyping |
| *Sm22-Cre* Rev | CCATGAGTGAACGAACCTGGTCG | Genotyping |
| *iMYOCD.* Fwd | ATCTGCAACTCCAGTCTTTCTA | Genotyping |
| *iMYOCD Rev* | CATATATGGGCTATGAACTAATGACC | Genotyping |
| 14,244,000-14,245,000 (-) | CATTAGGGTGTTAACAGTTTAGG | crRNA for Mapping *Myh11-CreER^T2^* |
| 14,276,000-14,277,000 (-) | TCCCAGTGTTCCCGCCAGGCAGG | crRNA for Mapping *Myh11-CreER^T2^* |
| 14,419,000-14,420,000 (+) | TGATTGACACCATTCTTGCTTGG | crRNA for Mapping *Myh11-CreER^T2^* |
| 14,270,000-14,271,000 (+) | CAGGGGTCCTAACCTTATATAGG | crRNA for Mapping *Myh11-CreER^T2^* |
| 14,307,000-14,308,000 (+) | ACGATTCCTTCATATGTGGAAGG | crRNA for Mapping *Myh11-CreER^T2^* |
| 14,318,000-14,319,000 (+) | ATCTTGTAGTCCACGCTATCTGG | crRNA for Mapping *Myh11-CreER^T2^* |

**Antibodies**

| **Target antigen** | **Vendor or Source** | **Catalog #** | **Working concentration** | **Lot # (preferred but not required** |
| --- | --- | --- | --- | --- |
| SRF | Proteintech | 16821-1-Ap | 1:200 (IHC) 1:1000 (WB) | NA |
| GAPDH | Millipore | MAB374 | 1:5000 (WB) | 2470405 |
| ITGA8 | Santa Cruz | sc-365798 | 1:1000 (WB) | G2417 |
| SM-Cy3 Actin | Sigma-Aldrich | C6198 | 1:200 (IHC) | 058M4761V |
| Alexa Fluor 488 anti-rabbit | Life tech | A11008 | 1:200 (IHC) | 2018309 |
| Cre recombinase | Cell Signaling | 15036S | 1:1000 (WB) | 1 |
| Vinculin | GeneTex | GTX109749 | 1:1000 (WB) | 40037 |
| HA-Tag | Cell Signaling | 3724S | 1:1000 (WB) | Lot 8 |
| Alpha Tubulin | Millipore Sigma | T5168 | 1:1000 (WB) | 105483 |
| SRF | Abcam | ab252868 | 1:1000 (WB) | GR3348906-2 |

**Cultured Cells**

| **Name** | **Vendor or Source** | **Sex (F, M, or unknown)** |
| --- | --- | --- |
| MOVAS | ATCC #CRL-2797 | unknown |
| HCASM | Thermo #C0175C | unknown |
| Primary mouse aortic smooth muscle cell | Primary cultures in our laboratory | M |
| mESC 129S6 | Sigma-Aldrich # |  |
